# A Photonic Biosensor-Integrated Tissue Chip Platform for Real-Time Sensing of Lung Epithelial Inflammatory Markers

**DOI:** 10.1101/2022.06.21.497028

**Authors:** John S. Cognetti, Matthew T. Moen, Matthew G. Brewer, Michael R. Bryan, Joshua D. Tice, James L. McGrath, Benjamin L. Miller

## Abstract

Tissue chip (TC) devices seek to mimic human physiology on a small scale. They are intended to improve upon animal models in terms of reproducibility and human relevance, at a lower monetary and ethical cost. Virtually all TC systems are analyzed at an endpoint, leading to widespread recognition that new methods are needed to enable sensing of specific biomolecules in real time, as they are being produced by the cells. To address this need, we incorporated photonic biosensors for inflammatory cytokines into a model TC. Human bronchial epithelial cells seeded in a microfluidic device were stimulated with lipopolysaccharide, and the cytokines secreted in response sensed in real time. Sensing analyte transport through the TC in response to disruption of tissue barrier was also demonstrated. This work demonstrates the first application of photonic sensors to a human TC device, and will enable new applications in drug development and disease modeling.

## Introduction

Currently, animal studies represent the gold standard for research on disease pathophysiology as well as for preclinical drug studies. While animals have appropriately complex physiology on an organ level, these studies often fail to translate to humans due to the genetic differences between the animal species being used and humans^1^. Animal research is also labor-intensive and expensive, and incurs high ethical costs due to the large number of animals used to obtain statistical significance. In a similar vein, clinical studies done in humans contain the maximum level of complexity, and have obvious relevance to humans. However, the vast heterogeneity between individuals in a sample population can distort the results^2^, and very large sample sizes are needed to prove marginal effects.

In contrast, *in vitro* models are able to remove the heterogeneity confounding human and animal studies, and are less expensive and less labor-intensive than animal studies. However, early *in vitro* studies lacked complexity, and thus yielded no information on the complex interactions occurring in tissues *in vivo*. This has led researchers to develop tissue chip (TC) platforms to study diseases and drug response^3,4^. These TC models greatly reduce reagent cost by incorporating microfluidic delivery systems, and often use multiple cell types, sometimes employing a three-dimensional architecture, to mimic the complexity of a human organ system. This allows the study of disease and drug mechanisms with far greater fidelity than prior *in vitro* models, while removing the extraneous genetic heterogeneity and lack of translation from animal and clinical studies. While genetic heterogeneity is a difficult reality of developing therapeutics, using genetically homogeneous sources (such as primary human cells or cell lines) can be useful for isolating basic mechanisms by removing the upstream or downstream factors that can be confounded by the complex gene pools involved in clinical trials. Additionally, the use of single-donor induced pluripotent stem cells as the starting point for TCs is anticipated to help identify patient-specific pathways in disease and therapeutic studies^5,6^.

A key limitation with TC studies reported to date is that there are very few methods available to determine real-time responses of cells to stimuli *in situ.* Existing TC models are able to measure various parameters including media oxygenation^7–9^, pH^10,11^, or glucose content^12,13^. Some use fluorescent markers or fluorescently-labeled antibodies to measure permeability of tissue barriers^14,15^ or track specific analytes, sometimes in separate endpoint assays^16^. However, these models rely on the availability of fluorescent tags, and their addition risks disturbing the system.

Recently, a few models have incorporated antibody-based electrochemical sensors serially in microfluidic tissue chips^17,18^. These models represent an important first step, but incorporation of the sensors relatively far from the cellular source of the analytes being measured is less than ideal. There is thus a recognized need in the field for methods enabling the incorporation of on-board, label-free sensors to measure the real-time dynamics of cellular interactions^19^. Time-resolved data on cellular secretion would yield precise information on drug and protein interactions with different cell types, enhancing the efficiency and accuracy of therapeutic trials.

Additionally, barrier permeability is an important marker of dysfunction in many diseases. To that end, some tissue chip models incorporate transendothelial electrical resistance (TEER) measurements^20,21^. However, TEER has recognized limitations in its ability to determine overall barrier integrity in a model device, as even a small hole in the monolayer will result in greatly reduced TEER values, even when most of the barrier has fully formed tight junctions. Some models have attempted to address this by either using movable electrodes^22^ or electrode arrays^23^, but still lack specific biosensing capabilities.

Here, we demonstrate the use of photonic biosensors to measure the secretion of cytokines from cells in response to a stimulus, and to detect an exogenous analyte passing through a disrupted tissue barrier. Realtime detection of cytokine secretion is a goal for TCs because cytokines are an important parameter in many inflammatory disease models, such as lung infection^24,25^ (akin to the model studied here), Alzheimer’s Disease^26,27^, and atopic dermatitis^28,29^. Additionally, such pathophysiology has already been studied in TC models or *in vitro* endpoint assay experiments. As mentioned above, earlier models have biosensors arranged as serial devices in a multi-device fluidic system. This dilutes the sample considerably as much of the analyte is lost to the device walls and tubing that connects the devices. Thus, the driving design parameter here was to incorporate the photonic biosensor in extreme proximity to the tissue culture substrate in order to maximize sensor performance. The device involves two microfluidic channels separated by a thin nanoporous membrane serving as the substrate for the culture of one or multiple cell types. A photonic biosensor chip is situated below the bottom channel. Analytes secreted by the cells or analytes introduced into the top channel moving through a disrupted barrier diffuse to the sensor chip, which is able to quantify the response in real time without any labels. A schematic of this platform is shown in Figure 1a.

**Figure 1.**
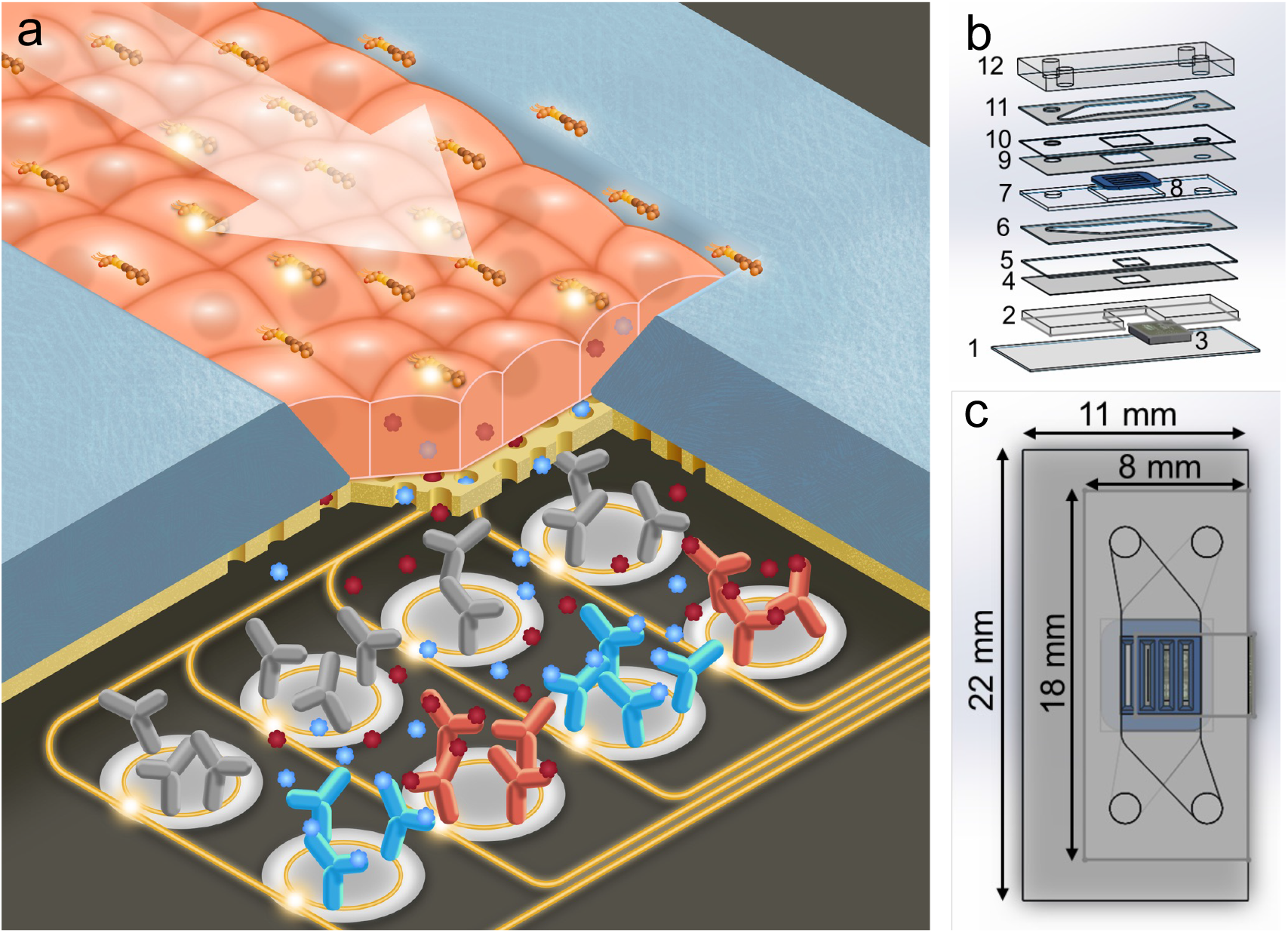
Photonic Sensor-Enabled Tissue Chip. a) Schematic of the working principle of the device. b) Exploded view with layers: 1) 170 μm glass coverslip, 2) 762 μm photonic chip holder, 3) photonic chip, 4) 57 μm adhesive sealing layer, 5) 127 μm silicone sealing layer, 6) 127 μm adhesive bottom channel, 7) 300 μm membrane chip holder, 8) four-slot nanoporous membrane chip, 9) 57 μm adhesive membrane sealing layer, 10) 100 μm silicone sealing layer, 11) 127 μm adhesive top channel, and 12) ~2 mm PDMS cap with inlets and outlets. c) Top view with outer dimensions of the device.

Photonic biosensors, and in particular devices based on photonic ring resonators, have been studied extensively due to their label-free nature and high sensitivity^30,31^. Ring resonator biosensors have also been commercialized (Genalyte, San Diego, CA; SiPhox, Burlington, MA, among others). Typically fabricated using standard CMOS processes in silicon or silicon nitride, ring resonators work by coupling light from a bus waveguide into adjacent rings producing resonances at certain wavelengths, according to the geometry of the ring and the effective refractive index of the medium in which the ring resonator sits. Since the electromagnetic field is not completely confined to the waveguides, it creates an evanescent field which decays in strength as a function of distance from the waveguide surface. This means that the resonant wavelength is affected by the refractive index of materials in close proximity (~100 nm) of the waveguide. Thus, binding of high-refractive index materials (e.g., proteins) to antibody-functionalized waveguides will result in a shift of the resonant wavelength toward the red end of the spectrum in a manner that can be calibrated to provide a quantitative readout as a function of analyte concentration.

Resonance-based photonic sensors have been shown to have very high sensitivity when operating in label-free mode, and signal enhancement via flowing a sandwich antibody or other strategies can push this to the pg/mL level^32,33^. However, the detection limits needed for a tissue chip platform are unclear. Clinical studies measure serum concentrations at single pg/mL or below^34,35^. Dilution into the serum means that this represents only a small fraction of whatever analytes are being produced in the tissue parenchyma of interest, and the concentrations of analyte in close proximity to the tissue are likely to be much higher. A recent study found single-cell secretion of cytokines to be in the ng/mL range within a few tens of microns of the cell^36^, but levels near fully developed tissue cultures or TC systems are unknown, and represent an important target of study and further motivation for our work. In collaboration with AIM Photonics in Albany, NY, we have developed designs and manufacturing processes for silicon nitride photonic ring resonators, and have validated their use as biosensors^30,37,38^. Sensor evaluation relied on a pressure-driven microfluidic platform for precise delivery of analytes, using a craft cutter to cut layers of silicone and adhesive tape, and bonding them with UV/ozone treatment, described previously^39^. Results from this system suggested that an analogous device incorporating an *in vitro* tissue culture could be suitable for measuring the secretion kinetics and concentrations of analytes in close proximity to the TC, yielding a better understanding of the dynamics of disease pathophysiology.

In order to integrate photonic sensors with cell culture, we developed a two-channel microfluidic design, with the two channels flanking the apical and basal faces of the model barrier. Media flows in the top channel, and cellular secretion can be monitored by the photonic sensor integrated in the bottom channel (Figure 1 b,c). As a substrate for culturing cells, we utilized nanoporous silicon nitride (NPN) membranes, a well-established platform for microfluidic applications^40^ commercialized by SiMPore, Inc. (West Henrietta, NY). These membranes are less than 100 nm thick, allowing for free diffusion of biomolecules which are much smaller than the ~ 30 nm pores.

For these experiments, bronchial epithelial cells of the 16HBE cell line were used, as they readily form tight monolayers *in vitro*^41,42^. We have previously studied these cells *in vitro* in Transwells™ as a model barrier system for testing tight junction-disrupting peptides (TJDPs), and have verified that they produce a robust network of tight junctions^43^. 16HBEs have been shown to secrete inflammatory cytokines in response to stimulation with the bacterial endotoxin lipopolysaccharide (LPS), with an explicit timecourse^44–46^. As a proof-of-concept, we sought to replicate this time course in our sensor-enabled microfluidic device by continuously flowing LPS over a monolayer of HBE cells, and measuring the sensor response continuously over the course of several hours, with a photonic sensor chip functionalized with antibodies for IL-1β, IL-6, and CRP. Additionally, we demonstrated the utility of this platform for monitoring barrier integrity, by flowing CRP in the top channel and measuring it in the bottom channel before and after TJDP-mediated barrier disruption.

## Methods

### Photonic Sensor Chip Design, Fabrication, and Functionalization

Photonic ring resonator chips consisted of two banks of 14 ring resonators each. Each bank has an edge-coupled 8-waveguide array, with one input waveguide splitting into seven waveguides, and two ring resonators on each waveguide. The overall chip footprint is 5.8 x 4.0 mm, with the rings in a 7 x 2 ring array with 350 μm pitch in both directions (Figure 2a). The rings on each waveguide were designed to resonate around 1550 and 1551 nm, with a free spectral range of 2.2 nm. Waveguide dimensions and coupling gaps were optimized based on simulation and previous work^30,37^ to provide the sharpest possible resonance peaks (i.e. highest possible quality factor, or Q) at these wavelengths, as narrow resonances allow for higher 31,47 sensitivity measurements.

**Figure 2.**
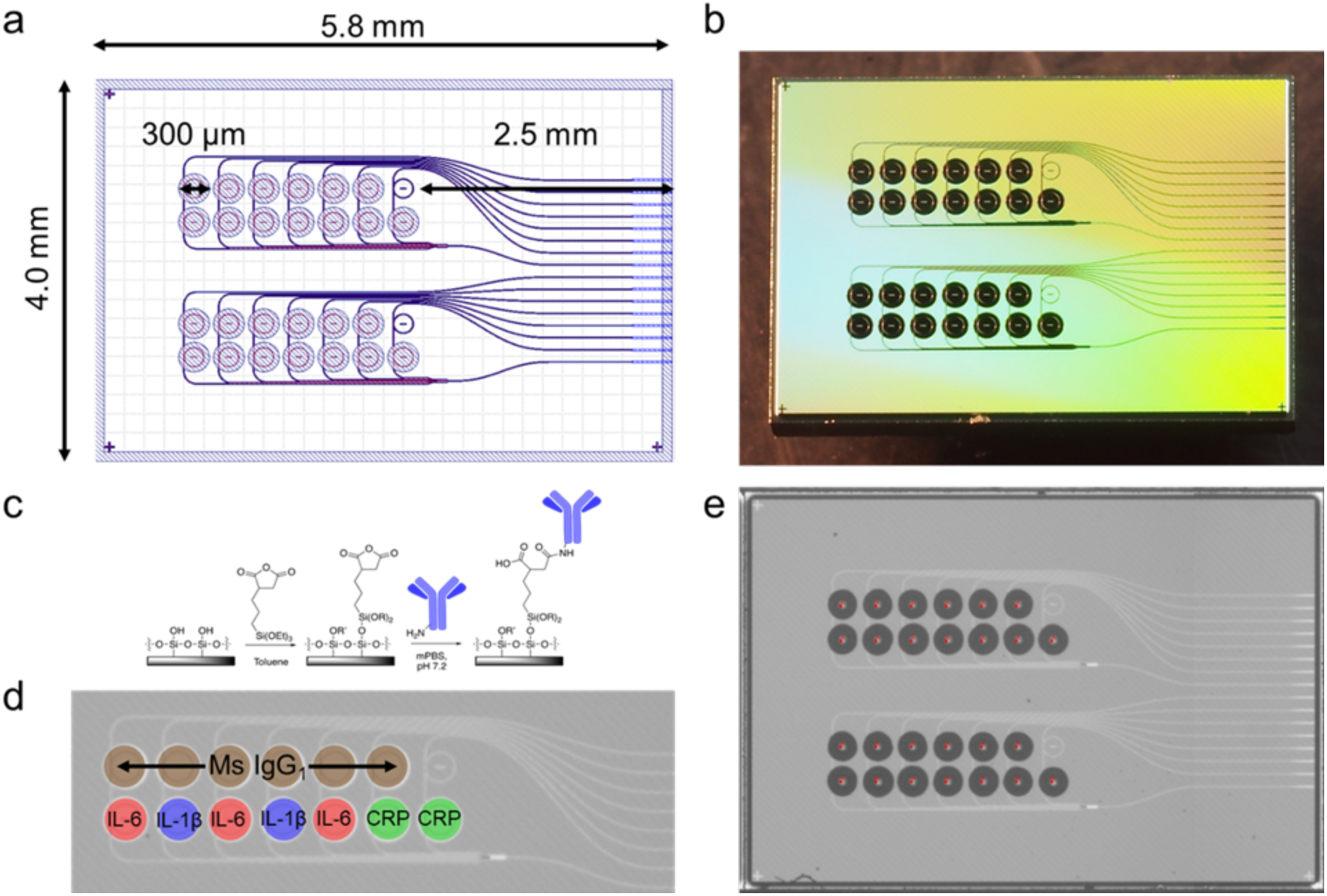
Photonic Chip Layout. a) Design in GDS (Graphic Design System) format used to produce photolithographic masks for the photonic chip. The overall footprint is 5.8 x 4.0 mm, with 2 banks of 7 x 2 ring resonators, each with a total diameter of 300 μm including the trench (open area in the top layer of SiO2 allowing direct contact between the ring resonator and analyte-containing solution). One temperature control ring (under oxide) is also provided in each bank. b) A photograph of the chip after fabrication and dicing. c) Functionalization chemistry using a succinic anhydride-bearing silane. d) Photomicrograph of a ring bank, with the antibody layout superimposed. e) An image acquired by the microarrayer immediately after spotting shows reproducible spotting of antibody solutions and StabilGuard over the rings. The dark spots are ~250 pL drops settled in the ring trenches.

Photonic sensor chips were fabricated using the 300-mm AIM Photonics fabrication line in Albany, NY that has been previously described^48^, using proprietary photolithography protocols. Briefly, about 220 nm of silicon nitride is deposited on top of 5.3 μm of oxide and patterned to form the waveguides. A subsequent 5-μm oxide layer is deposited on top. This oxide layer is then patterned with photoresist and selectively etched, to reveal the ring resonators. The 5 μm-deep trenches surrounding the rings are accessible to spotting of specific antibodies or analytes directly on individual ring waveguides.

Prior to functionalization, chips were cleaned with a 1:1 mixture of methanol and concentrated aqueous hydrochloric acid, rinsed three times with Nanopure water, and dried with a stream of nitrogen gas. They were then silanized via incubation in a 1% solution of 3-(triethoxysilyl)propyl succinic anhydride (Gelest, Morrisville, PA) in anhydrous toluene (distilled from Na metal under N_2_ atmosphere), rinsed for 5 minutes in pure anhydrous toluene, dried with N2 gas, and incubated at 110 °C for 30 minutes to allow the functional layer to completely anneal and evaporate any residual toluene. The functional chemistry is visualized in Figure 2c.

After silanization, antibody solutions were printed on the rings using a Scienion SX SciFLEXarrayer piezoelectric microarrayer equipped with a PDC-60 capillary nozzle. The microarrayer produces droplets of a particular size depending on the diameter of the capillary nozzle and the electrical settings on the piezoelectric controller. Here, droplets of about 250 pL each were dispensed, and the number of droplets was chosen to yield about 3 nL total spotted on each ring. This results in the antibody solution filling the trench around each ring without spilling to the neighboring trenches. In these experiments, the top ring (as viewed in Figure 2d) on each waveguide was spotted with mouse IgG_1_ isotype control antibody (R&D Systems, Minneapolis, MN) at 1000 μg/mL. Mouse IgG_1_ isotype control antibodies were chosen to ideally match the nonspecific binding characteristics of the capture antibodies we used, which were all of the mouse IgG_1_ isotype. Subtracting the response of the control rings from the capture rings yields the response due only to specific binding of antibody/antigen pairs. The bottom rings were spotted with antibodies to IL-1β (R&D Systems) at 500 μg/mL, IL-6 (Biolegend, San Diego, CA) at 650 μg/mL, IL-8 (Sino Biological, Wayne, PA) at 1000 μg/mL, and C-reactive protein (CRP; Ortho-Clinical Diagnostics, Rochester, NY) at 990 μg/mL (Figure 2d). When dilution was necessary from the stock concentration, antibodies were diluted with PBS at pH 5.8. After allowing the antibodies to covalently attach to the functionalized rings for 30 minutes, StabilGuard (Surmodics, Inc., Edin Prairie, MN) was overspotted on the rings as a stabilizer, keeping the chips shelf-stable until they were ready to be incorporated into a device and used (Figure 2e).

### Nanoporous Silicon Nitride (NPN) Membrane Cell Culture Substrates

NPN membrane chips were purchased from SiMPore, Inc. (West Henrietta, NY). Nanoscale porous membranes were chosen for this research because their thinness allows for free diffusion of analytes, in contrast to popular polymer track-etched membranes, which are about 10 μm thick. NPN membranes from SiMPore are approximately 70 nm thick, which also allows for acquisition of high-quality microscopy images of cells cultured on the membrane. This 70-nm layer of nitride is deposited on a silicon wafer, and the silicon is then etched away from the back to reveal the porous nitride membrane.

NPN membranes have been demonstrated to be excellent cell culture substrates^49,50^. Additionally, their thinness means that they are optically clear, enabling high-quality transmission microscopy. A four-slot/four-membrane design was chosen so that one membrane could be overhung beyond the silicon photonic sensor chip, allowing visualization of the cell monolayer via transmission-based phase contrast microscopy through the device, while the other three membranes are situated directly above the sensor chip. The membrane slots are 3 mm long by 300 μm wide, on a 5.4 × 5.4 mm silicon chip frame. The silicon is 310 μm thick. Due to anisotropic etching of the silicon, the trench opposite each membrane is wider than 300 μm, which needed to be accounted for when designing microfluidic sealing layers. Membrane chip dimensions are shown in Figure 3.

**Figure 3.**
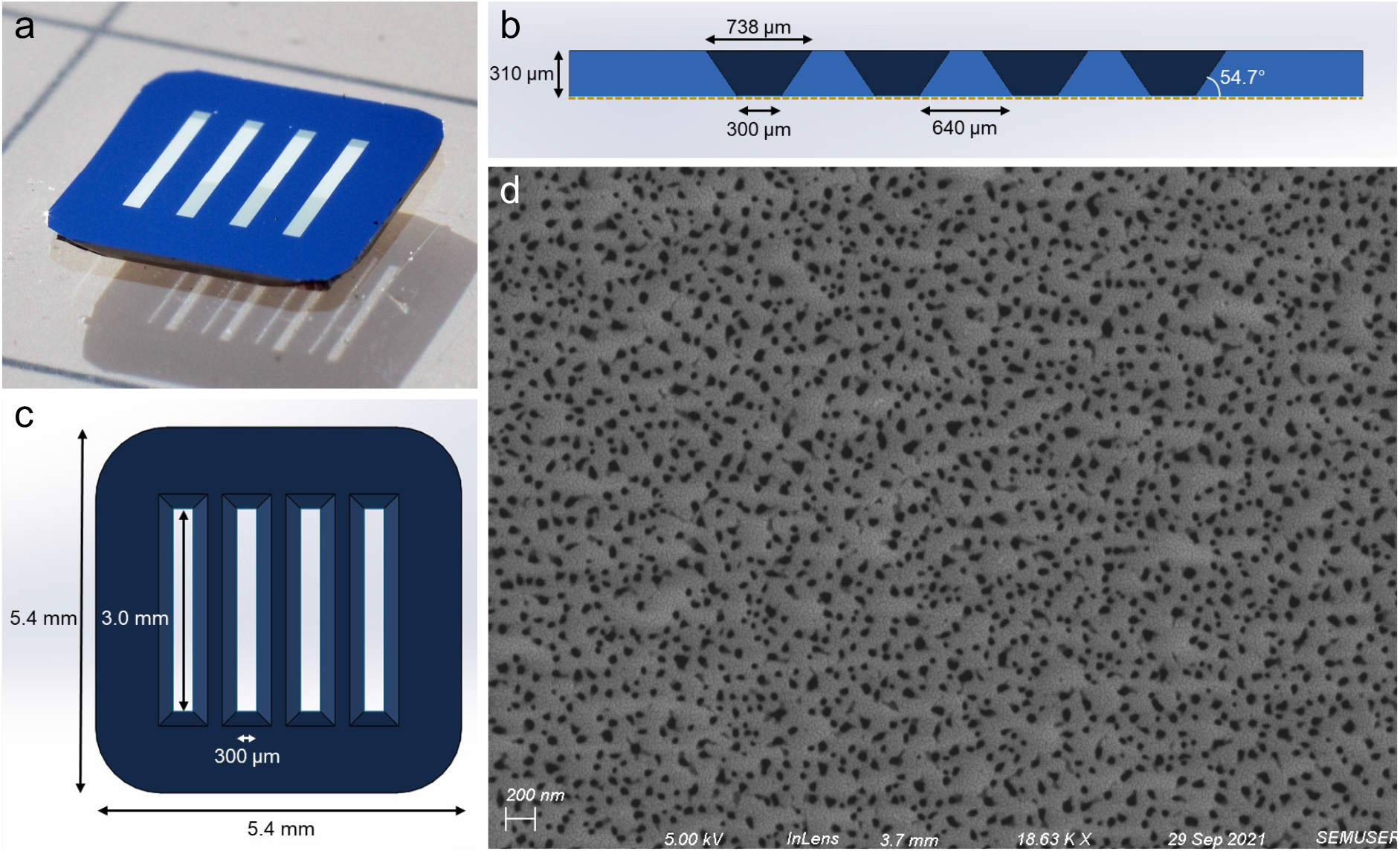
Membrane Chip Design. a) Photograph of a four-slot NPN membrane chip (color is due simply to lighting), with the etching trenches face down. b) Schematic cross section of the chip showing the anisotropically etched trenches opposite the nanoporous membrane (shown as a gold dotted line). c) Top view schematic of the 5.4 × 5.4 mm chip, with membrane slots that are 3.0 mm × 300 μm. d) An SEM image of the membrane, showing a heterogenous distribution of pores, averaging 28 nm in diameter. The membrane has an overall 13.5% porosity.

### Microfluidic Device Design and Fabrication

The devices used here required the incorporation of a membrane chip for cell culture sandwiched between two microfluidic channels, with a photonic chip incorporated at the base of the bottom channel. This introduced a number of constraints that influenced the device design. First, the multichannel design required a layered approach. Thus, layers of silicone and adhesive tape were cut using a commercial craft cutter (Cameo 3, Silhouette America, Lindon, UT) and stacked via UV/ozone bonding^51,52^ to both hold the membrane and photonic chips, and form channels. Also, due to the photonic chip requiring input from and output to an optical fiber array on the side of the chip, the photonic chip had to project from one side of the device, so that the edge coupling array was exposed. Another consideration was that due to the opacity of the silicon photonic chip, cells could not be visualized with brightfield or phase contrast microscopy. We therefore chose to place the membrane chip such that one of the four membranes overhangs the photonic chip, enabling monitoring of the cell culture by microscopy.

With these design constraints in mind, we designed a two-channel device stack to include a 170 μm thick glass coverslip (cut in half for a 22 x 11 mm footprint), 762 μm silicone (Specialty Manufacturing Inc., Saginaw, MI) to hold the photonic chip in place, then two identical sealing layers, first of 57 μm adhesive (3M, Maplewood, MN), then of 127 μm silicone (SMI). The membrane chip is encased in a layer made from 300 μm silicone (Trelleborg Sealing Solutions, Trelleborg, Sweden). The membrane layer is sealed with two more identical layers: 57 μm adhesive, and 100 μm silicone (Trelleborg). The top channel is also 127 μm adhesive. The device is finally closed with polydimethylsiloxane (PDMS, Sylgard 184, Dow-Corning, Corning, NY) with two inlet and two outlet holes, made with 21-gauge blunt dispensing tips (Jensen Global, Santa Barbara, CA). The device layer stack is seen in Figure 1b.

All silicone, glass, and PDMS was activated with UV/ozone (Novascan PSDP, Novascan Inc., Boone, IA) for 5 minutes prior to adhesion. Device layers were aligned manually with forceps and pressed firmly together to ensure a good seal. The membrane chip plus sealing layers were heated at 70 °C for about 5 minutes to ensure a good seal. Manual assembly of these devices took about an hour and a half, though multiple devices could be assembled in parallel to increase throughput.

For analyte calibration experiments, a single-channel device design was used. This device consists of the coverslip, 762 μm silicone chip holder with the photonic chip nested inside it, 57 μm adhesive seal, 127 μm silicone seal, 127 μm adhesive channel, and PDMS. These devices took about 15 minutes to assemble.

The single-channel device stack can be seen in Figure S1. For each assay, known concentrations (in factors of ten from 10 pg/mL to 10 μg/mL for CRP and to 1 μg/mL for the other analytes) were flowed over the chip for 25 minutes each, a sufficient time for equilibrium to be reached as determined by no further shift in sensor resonance occurring. Sensor responses (relative shift at each concentration) were fit to a four-parameter logistic curve using Origin Graphing Analysis Software (OriginLab Corp, Northampton, MA).

### Cell Culture

Human bronchial epithelial cells (16HBE line) were cultured in DMEM media (Gibco, Bleiswick, Netherlands), supplemented with 10% fetal bovine serum (Gibco), 1% HEPES (Gibco), and 1% penicillin/streptomycin. Cells were cultured in T25 flasks to confluency, which usually took 5-7 days, and passaged.

Prior to seeding cells in a device, the membranes were coated with extracellular matrix proteins to facilitate cell adhesion. To do this, silicone tubing with a 0.02” inner diameter (Cole-Parmer, Vernon Hills, IL) was inserted into the inlets and outlets with a 21-gauge 90° cannula (Jensen Global). The bottom channel was injected with PBS (pH 7.4), and the top channel was filled with a mixture of rat tail type I collagen (Sigma Aldrich, St. Louis, MO) at about 100 μg/mL, and human fibronectin (Gibco) at 50 μg/mL in PBS, and the tubing sealed with binder clips. The device was placed in a custom 3D-printed holder to hold the tubing in place while moving it, as tension in the tubes could cause them to detach or damage the device. It was then moved to an incubator for 2 hours. Media was then injected into the device to wash out any unattached matrix proteins. Meanwhile, cells from one T25 flask were pelleted, then reconstituted in 700 μL of media. This yielded an appropriate density for the cells to achieve a monolayer on the membrane quickly. Once cells were injected into the top channel, they were moved to the incubator and allowed to adhere to the membrane without flow for about 3 hours. At this point the device was taken out of the incubator, and a cannula was inserted between the top channel outlet and the bottom channel inlet, so that media could circulate continuously through both channels. A syringe was used to manually push out any bubbles that had formed. The device was then attached to a peristaltic pump (P-1, Cytiva, discontinued) in the incubator, with care to avoid introducing bubbles into the tubing.

The peristaltic pump was connected through the same 0.02” silicone tubing to two fluid reservoirs made from glass vials with 21-gauge cannula inlets sealed with silicone adhesive, which in addition to preventing bubbles from entering the tubing going to the device, also dampened the peristaltic flow from the pump, as previously described^50^. The pump was then turned on at the lowest setting, corresponding to about 30 μL/min, and left overnight for an experiment to be done the next day. This is because in our experience, sustained levels of high shear stress result in poor cell health.

For long-term cultures under such low flow rates, bubbles spontaneously form over the membranes. By covering the device in the incubator with a 3D-printed enclosure, with two 15-mL conical tube caps filled with distilled water, the local humidity was increased to a level that prevented and eliminated existing bubbles completely over the course of all long-term incubations. The incubator setup is shown in Figure 6a.

**Figure 4.**
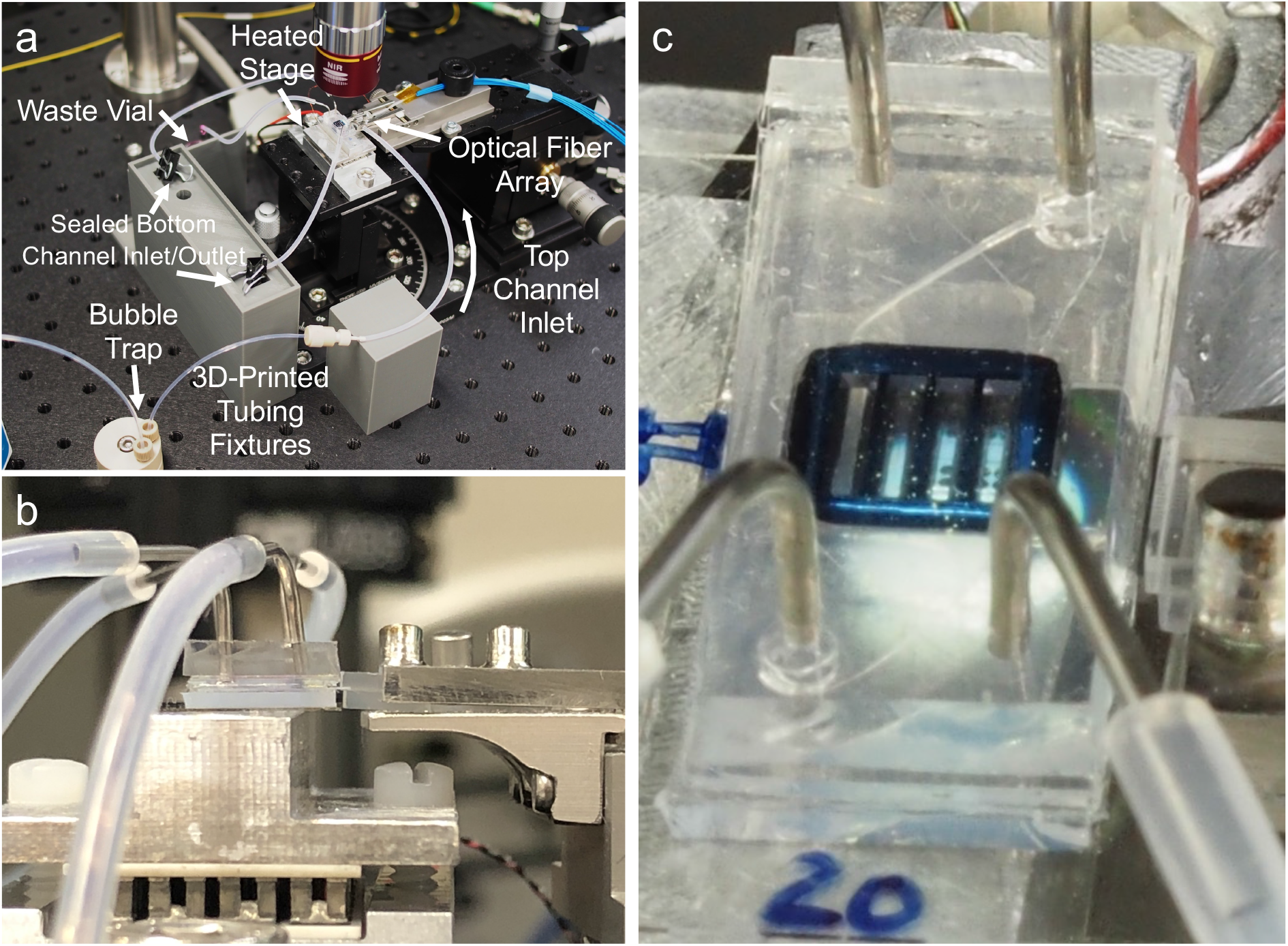
Optical Configuration. a) The device is situated on a heated aluminum stage to be addressed by a fiber array. Fluidic tubing for both channels is held in place by custom 3D-printed fixtures. The bottom channel remains sealed for the duration of the experiment. b) A side view of the fiber array approaching the photonic chip from the right side with the device on a heated aluminum stage. c) A closeup of the device and fiber array, with rings under the membranes, photonic waveguides, and optical fibers visible.

**Figure 5.**
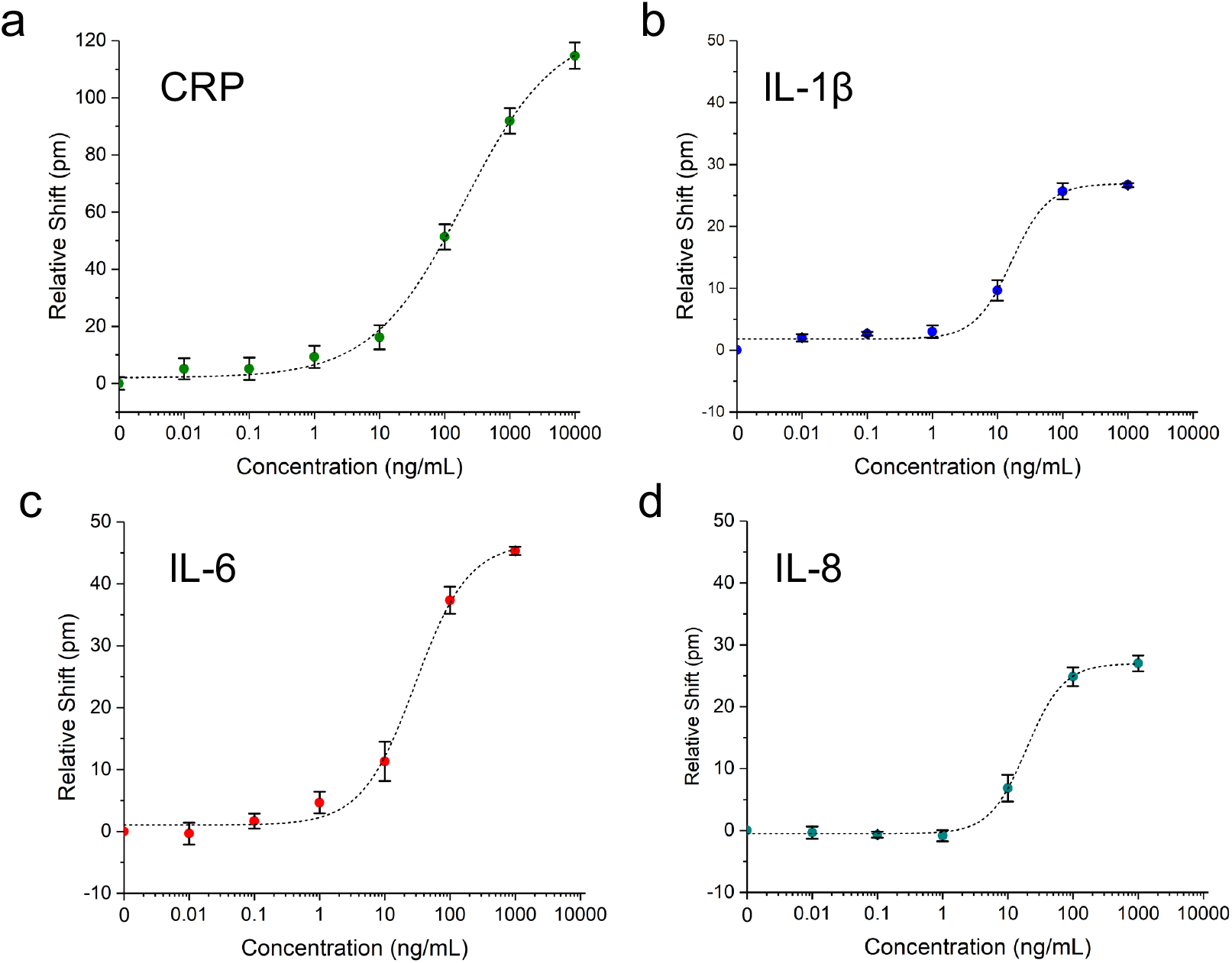
Photonic sensor validation in a single-channel microfluidic device. Photonic sensor calibration for various analytes: a) CRP (n = 6 channels), b) IL-1β (n = 3), c) IL-6 (n = 3) and d) IL-8 (n = 4). All responses were fit with four-parameter logistic curves. Error bars represent standard error of the mean.

**Figure 6.**
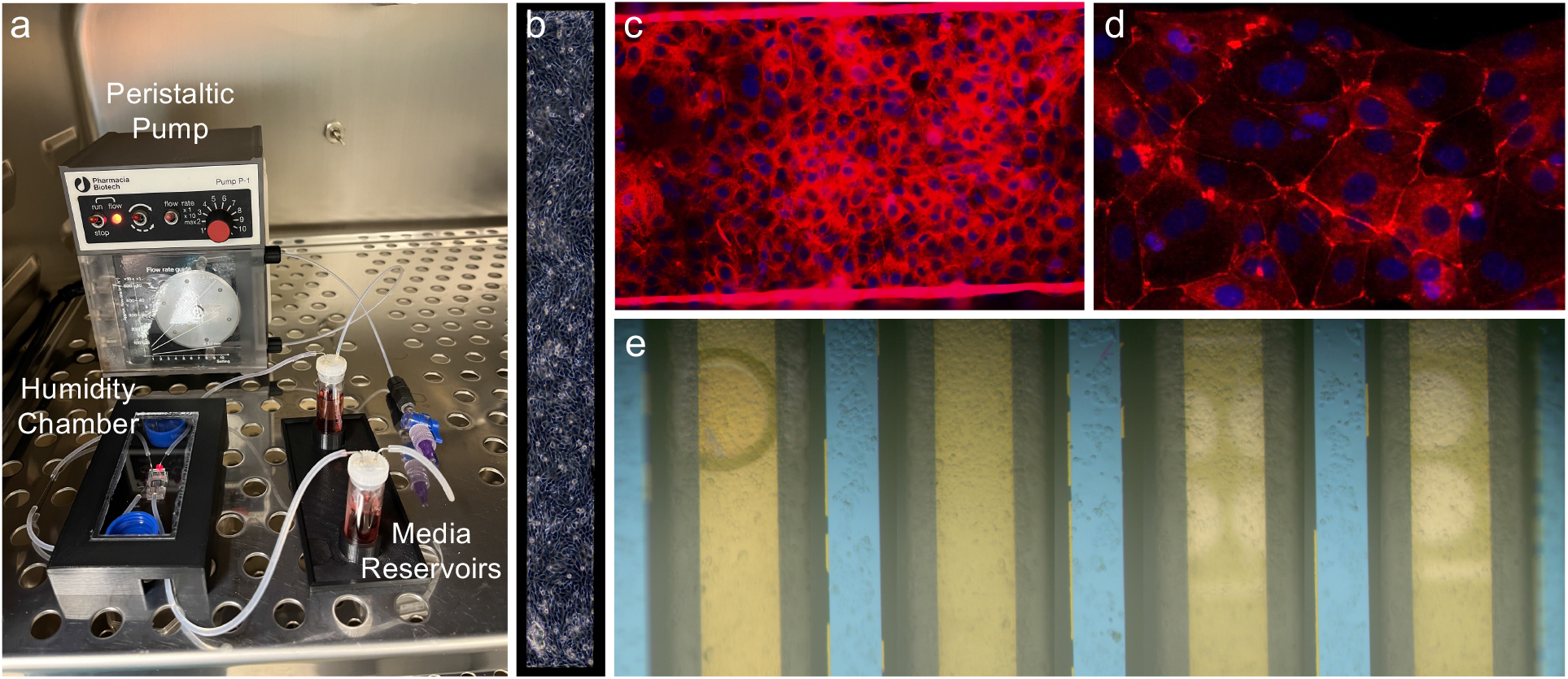
Cell Culture. a) Incubator pump/device interface. b) Phase contrast image of the overhung membrane with a monolayer of HBEs. c) Tight junction staining of HBEs on a membrane (the bright lines represent cells growing up the edge of the membrane suspended between edges of the etched silicon substrate) (20x) and d) the silicon supports between the suspended membranes. (40x) e) Stitched brightfield image of all four membranes with cellular monolayers prior to a sensing experiment. The photonic ring resonator trenches (white) are visible underneath the two right-most membranes. While bubbles over or in close proximity to the ring resonators alter their resonance wavelengths, the bubble seen on the fourth (leftmost) membrane here will not interfere with cell secretion measurements as it is far away from the sensors.

### Optical Configuration

To acquire resonance shift information from the sensors in real time, the microfluidic device with integrated photonic sensor needed to be coupled to a light source and detector. To accomplish this, the device was situated on an aluminum stage fitted with a thermoelectric coupler (TEC) and thermistor to maintain the device at 37 °C. Light from a tunable laser source (Keysight 81606A) was guided through a polarization controller via an FC/APC single-mode fiber (Thorlabs Inc., Newton, NJ) and approximately polarized with TE orientation relative to the rectangular waveguides of the photonic chip. It was then routed into the first fiber of an 8-fiber linear fiber array with a 127-μm pitch (Oz Optics, Rochester, NY). The 7 output fibers were attached to an optical power meter (Keysight N7745C). The 8-fiber array was then aligned to the 1-input-7-output waveguide array on the right edge of the photonic chip, visualizing with both a visible camera (Thorlabs DCC1645C) and an IR camera (WiDy InGaAs 650) and adjusting with micrometers, until the power registered in the output channels is maximized. Though there are two banks of rings on these chips, only one was used for these experiments due to the use of 8-channel detectors. In the future, more ring banks can be utilized by the use of detectors with more channels or by configuring multiple detectors.

Fluid delivery was controlled with a pressure driven fluidic pump (Aria Cell Perfusion System, Fluigent Inc., Le Kremlin-Bicêtre, France), that can be preloaded with up to ten different samples. A switch allowed seamless transition between samples, and a bubble trap prevented air from disrupting the cells in the device.

### Data Analysis

To calculate the resonance shift due to binding, resonance peak locations had to be extracted from the raw data. A custom Python script converts Keyence .omr spectral files into a single .parquet file, described previously^37^. Then a user interface allows the user to choose an experiment and channel, and then finds and plots peak locations across the entire experiment (Figure S2). The user manually assigns peaks to either the “control” or “capture” rings, and the script subtracts the control shift from the capture shift. The resulting “relative shift” represents the shift in the experimental rings due only to the capture of specific analytes. These values are all exported as .csv files.

### Immunocytochemistry

To confirm the ability of HBE cells to form a robust barrier in the device, cells were cultured in a device for four days under minimum flow conditions (for the peristaltic pump used, the minimum was about 30 μL/min), and then stained for ZO-1, a common marker of tight junctions^53^. Fixation and staining were done in the closed device by injection of reagents through the tubing with a syringe. On the fourth day cells were fixed with 4% paraformaldehyde at room temperature for 1 hour, then methanol at −20 °C for 20 minutes. After washing 3x with PBS (pH 7.4), cells were blocked with 1% bovine serum albumen (BSA) in PBS. Then primary antibody, mouse anti-ZO-1 (Invitrogen, Waltham, MA), diluted 1:300 in 1% BSA solution, was added and incubated at room temperature for 2.5 hours. After 3 more rinses with PBS, secondary antibody, Alexa-568-conjugated donkey anti-mouse (Life Technologies, Carlsbad, CA) diluted 1:1000 in 1% BSA, plus DAPI (Molecular Probes, Eugene, OR) diluted 1:1000, was added and incubated at room temperature for 1.5 hours. After three more rinses in PBS the entire device was imaged via fluorescence microscopy (Olympus BX-60) with PBS still in the channels.

### Cytokine Sensing Experiments Following LPS Stimulation

Sensing experiments were conducted one day after seeding cells in a device, and had a full or near-monolayer of cells covering all four membrane slots. Fluid reservoirs of the Fluigent Aria were pre-filled with all of the solutions to be used. The cell- and sensor-containing device was adhered to the aluminum optical stage with adhesive, and the TEC element was set to 37 °C. In the incubator, media was recirculated from the top channel outlet to the bottom channel inlet. However, for sensing experiments, the two channels were disconnected from each other and the bottom channel was sealed with binder clips in order to allow for analyte capture by passive diffusion from the cells. The top channel inlet tubing of the device was connected to the outlet tubing of the Fluigent pump, and media was flowed at 30 μL/min continuously through the top channel. The photonic chip was then aligned to the fiber array, and spectral measurements then began immediately (typically within 5 minutes of placing the device on the optical stage). The laser was set to scan a 6-nm wavelength range (1547-1553 nm) at 1 pm resolution at 19-second intervals continuously throughout the duration of the experiment.

Occasionally, as is typical with microfluidics experiments, a bubble would appear in the top channel, in which case flow would be briefly disconnected to force the bubble out with a media-filled syringe. Then the chip would be realigned to the fiber array, if necessary. This usually only manifested as a transient bump in the spectral peaks, but for both experimental and control rings equally, thus not affecting outcomes. However, a bubble in the bottom channel was more problematic, as a low-refractive index mass nonuniformly distributed over the rings drastically altered spectral features. Therefore, any experiments with significant bubble interference in the bottom channel were unusable, and that data was discarded.

### Sensing of Analytes Through Disrupted Barrier

16HBE cells were seeded in a device and allowed to form a tight barrier under minimal flow conditions −(30 μL/min). Cells had formed a full monolayer after 1 day. After 4 days, the barrier was presumed to have formed tight junctions (based on the ZO-1 staining experiments described above), such as to prevent the passage of large analytes through the paracellular spaces. The device was moved to the optical setup and continuous spectra were taken, while first flowing media for 30 minutes to allow the sensors to equilibrate, followed by 1 hour of CRP at 1 μg/mL.

Before reattaching the device to the peristaltic pump, the media was supplemented with tight junction-disrupting peptides (TJDPs) at 10 μM (dosage demonstrated previously^43^), and the peptides were allowed to flow over the chip overnight. Approximately 22 hours later (day 5), the device was reattached and the again media was flowed for 30 minutes followed by CRP at 1 μg/mL for 1 hour. Then the pump inlet was switched to the bottom channel, to allow CRP to flow directly over the sensors. This was done as a positive control, which also yielded information about the relative amount of CRP that was able to diffuse through the disrupted barrier.

## Results

### Photonic Chip Sensitivity

Photonic ring resonator biosensors yield transmission spectra containing resonance troughs at the wavelength resonant for the effective refractive index of their environment. These chips contained waveguides with two ring resonators each, with one designed to have a resonance trough at 1550 nm and the other at 1551 nm, by altering the diameter of the rings by a minute amount. The rings each have a free spectral range of about 2.2 nm, leaving enough distance between repeating comb features to prevent the crossing of peaks for the shifts seen here. The rings used here have quality (Q) factors (a measure of the sharpness of the resonance troughs) above 10^5^, in line with current state-of-the-art for photonic biosensors^31^. An example set of spectra for all 7 channels of a representative sensor is shown in Figure S2a.

Using single-channel fluidic devices, sensor calibration experiments were performed for four analytes. C-reactive protein (CRP) was used as a positive control for sensor function, and a negative control for cell stimulation. While CRP is a common marker of inflammation, it is not produced by HBE cells^54^, and with a mass of 120 kDa, the CRP pentamer is easily sensed in a label-free assay. IL-1β (17.5 kDa) and IL-6 (21 kDa) are common pro-inflammatory cytokines, and were calibrated on the same multiplex chip, which had four waveguides functionalized for IL-6 and three for IL-1β. IL-8 (8.5 kDa) is another pro-inflammatory cytokine calibrated on its own chip.

For a high concentration of 1 μg/mL of analyte, the various protein markers yielded responses roughly proportionate to their molecular weight. CRP had a relative (i.e. control-subtracted) shift of 92 pm (n = 6 channels), IL-1β (n = 3), with IL-6 (n = 3), and IL-8 (n = 6) shifting 27, 45, and 27 pm, respectively. Measured lower limits of detection for these analytes were 1.5, 3.1, 7.6, and 20.7, for IL-1β, CRP, IL-6, and IL-8, respectively. However, as mentioned earlier, the actual concentration near tissue will likely be much higher. Additionally, the consistency of ring-to-ring response within a single chip allows for the rigorous quantification of secreted analytes in our cellular devices. While duplicate rings on a single chip produced nearly identical results, chip-to-chip and wafer-to-wafer variability has yet to be assessed yet, and will need to be examined further. We anticipated that these limits of detection would be sufficient for our planned experiments, but, as mentioned previously, sandwich assays and other strategies could be used to enhance signal as needed^33^.

### Cell Viability in Devices

To demonstrate that 16HBE cells behave as expected in the context of our microfluidic system, relative to other studies using microfluidic systems to culture HBEs^55,56^, we stained them for the tight junction marker ZO-1 after four days. The images shown in Figure 6 indicate the presence of tight junctions in cells growing both on the free-standing NPN membrane (Figure 6c) and on the silicon-backed regions between membranes (Figure 6d). The ZO-1 is diffuse in the membrane domains compared to tight junction staining seen on 16HBEs in literature, but it can be clearly visualized in the silicon-backed regions of the membrane chip (Figure 6d). Though experiments employing LPS stimulation did not require tight junctions, longer-term culture and an intact barrier were required for experiments testing analyte diffusion through a TJDP-disrupted barrier.

Visualizing cells via phase contrast microscopy (Figure 6b) or fluorescent staining (Figure 6c) was easily done via on the membrane overhanging the silicon photonic chip. However, cells on other membranes were also easily visualized with reflective brightfield microscopy on the optical setup. When performing a sensing experiment, cell morphology was monitored throughout the duration of the experiment by focusing on the membranes with the visible light camera, allowing the extent of the monolayer or loss of cells under flow to be observed (Figure 6e).

### Nonspecific Binding Control and Diffusion through Bottom Channel

To understand nonspecific binding over timescales relevant to cell sensing experiments, and to prove that analytes originating at the level of the membrane could reach the level of the sensors on a short timescale, an experiment was done with a two-channel, membrane- and sensor-containing device lacking cells. Media was flowed for three hours followed by 1 μg/mL of IL-1β, IL-6, and IL-8 in the top channel, in a similar manner to the single-channel calibration experiments. Despite the use of isotype-matched antibody controls, small differences in nonspecific binding between control and capture rings result in drifting relative shift over time when the shift of the control peak is subtracted from the shift of the capture peak. No drift in the temperature control ring was observed. However, as can be seen in Figure S3(b, d, f), there are slight linear signal drifts that differ for each analyte. These represent subtle differences in nonspecific binding between experimental and control antibodies, despite the use of class-matched controls. Importantly, the nonspecific shifts seen here are small enough that specific shifts due to binding of target analytes are readily observable. Once the analyte cocktail was added, relative shifts on par with single-channel experiments can be seen, with a 5-10 minute delay from when the analyte enters the top channel until it reaches the sensors at the floor of the bottom channel (Figure S3c, e, g). This highlights the potential for our system to acquire time-resolved data of cellular response to stimuli added to the top channel.

To develop a more detailed theoretical understanding of analyte and fluid flows in the device, a COMSOL simulation was run, mimicking the device’s geometry exactly (Figure S4a-d). A steady-state velocity profile (Figure S4e) shows that when the bottom channel is sealed, the membrane effectively isolates the bottom channel from flow in the top channel. This isolation is presumably enhanced with a cell monolayer cultured on the membrane. A separate simulation set the inflow concentration in the top channel at 1 μg/mL, and the diffusion coefficient set to 2.7×10^-7^ cm^2^/s to approximate that of IL-6 in solution^57^. A time-dependent simulation was then run to calculate analyte concentrations at 10-second intervals for up to ten minutes. The resulting simulation shows that the concentration at the level of the sensor reaches 50% of the input 1 μg/mL at about 90 seconds, and after 5 minutes about 95% of the input concentration. Concentration profiles and a graph of concentration at the level of the sensor are shown in Figure S5, and a GIF file of the concentration profile over the first 5 minutes is shown in Figure S6. This suggests that we can sense relevant quantities of secreted analytes on a timescale of minutes.

### Sensing of Cellular Secretions

As a proof-of-concept experiment, we tested the ability of our device to sense the secretion of cytokines from HBEs in response to stimulation with LPS. By flowing LPS in the top channel, cells are stimulated to produce and secrete inflammatory cytokines into the bottom channel. While the apical/basal secretion preference of 16HBE cells is unclear^58,59^, some other cell types have been shown to preferentially secrete cytokines basally^60^, i.e. opposite the side of stimulation. If the bottom channel is sealed, the secreted cytokines are allowed to diffuse to the level of the photonic sensor chip, situated at the bottom of the channel.

As seen in Figure 7, our experiments demonstrate the sensing of interleukins secreted in response to LPS stimulation, with a similar time course to previous experiments with HBEs. In the experiment shown here, media was flowed for 30 minutes, followed by LPS at 100 ng/mL for two hours. About one hour after the introduction of LPS, the sensor response, as represented by an increase in the relative shift of the IL-1β-or IL-6-functionalized rings, begins to increase steadily. After 90-120 minutes, the response levels off as the cell secretion level/analyte binding reaches equilibrium. The CRP-functionalized ring serves as an additional negative control (besides the Ms IgG_1_ control rings), and the trace shown in Figure 7c has no increasing trend like any of the cytokine-functionalized rings, confirming the specificity of the response to cytokines secreted by the HBEs. The slight negative drift for the CRP subtracted trace (and return to baseline drift after the deviation around 120-140 min.) likely represents a mismatch in the nonspecific response between the CRP antibody and the mouse control antibody, as discussed above.

**Figure 7.**
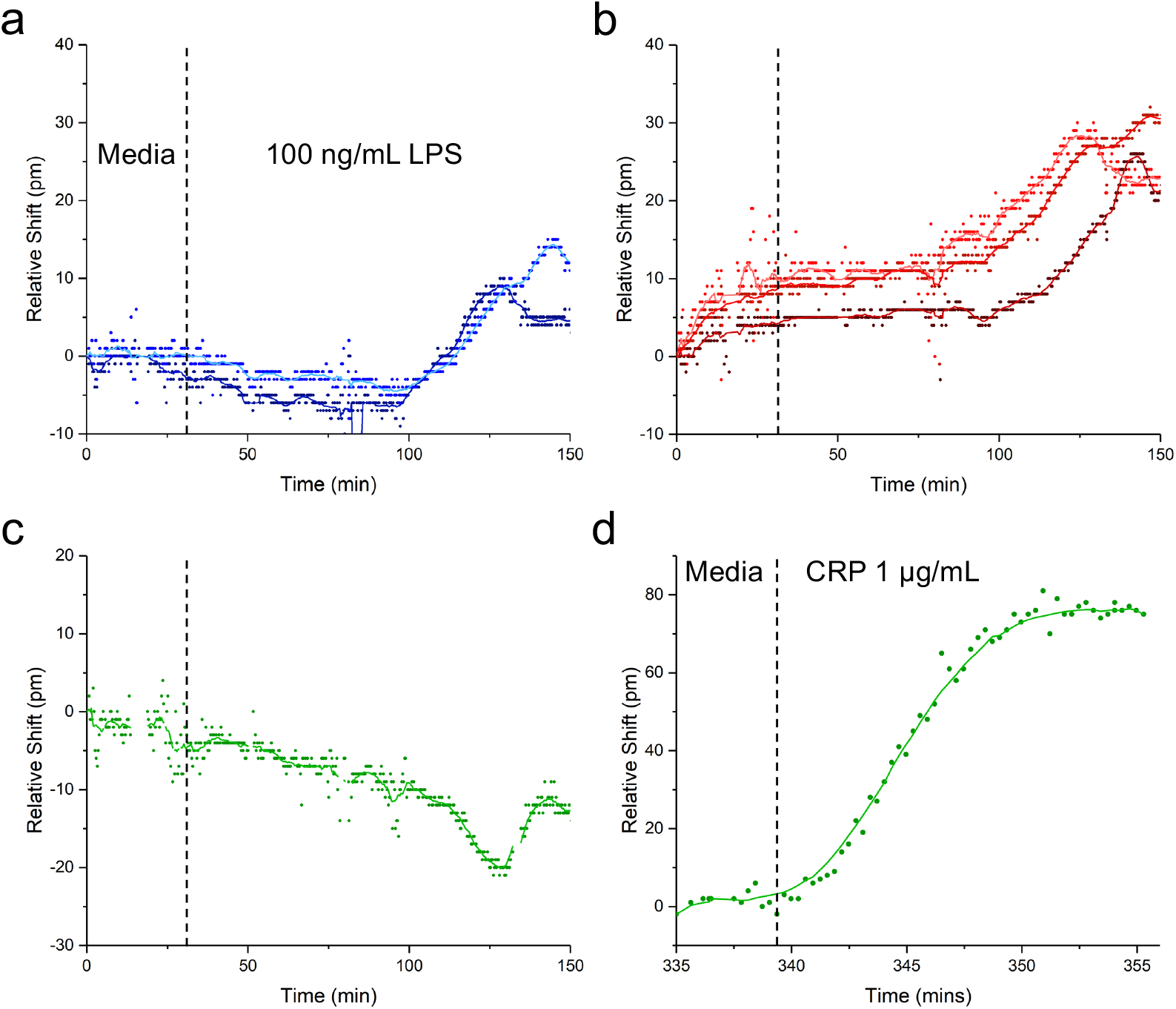
Sensing of Secreted Cytokines. Media flowed for 30 minutes before 100 ng/mL LPS was injected into the top channel and flowed for two hours. a) Response for two IL-1β channels, with the control response subtracted from the anti-IL-1β-functionalized ring response. The different shades of blue correspond to the individual IL-1β-functionalized channels. b) Subtracted response for three IL-6 channels. c) Subtracted response for CRP (i.e. negative control channel). d) The same CRP channel when recombinant CRP was added at the end of the experiment.

The CRP ring also serves as a positive control to verify that sensor responses seen are in fact due to antigens binding to antibodies bound to the rings, and that the sensor is still functional at the end of the experiment. In Figure 7d the trace for this ring is shown, in response to flowing 1 μg/mL of CRP in the bottom channel (i.e. directly over the rings) over the last 30 minutes of the experiment. There is a marked redshift in the resonance, with time-dependent binding response consistent with Langmuir binding behavior. The relative shift reaches a maximum of about 80 pm, compared with about 90 pm for the calibration experiment shown in Figure 5. None of the rings functionalized for other analytes showed this response (data not shown).

A separate experiment in which cells had peeled off the two right-most membranes prior to LPS stimulation provided an opportunity to test spatially-resolved sensing. LPS was flowed at 10 ng/mL and the sensor response was measured over the course of 4 hours. Figure S7 shows the responses across 6 channels, for IL-1 β (S7c) and IL-6 (S7d). The control-subtracted shift ranges from 1-15 pm, showing a decrease relative to the higher concentrations of LPS used earlier, as well as a temporal delay in the onset of response occurring at about three and a half hours. Additionally, there is an increased response with increasing channel number. Thus channel 7, which is the furthest channel to the left (closest to the remaining cells), had the highest response, with decreasing shifts for channels 6-4, until there was essentially no response in channels 3-1 (farthest away from the cells). Due to the loss of cells on the right side of the membrane chip, this demonstrates a graded response in accordance with the presence of full cell layers on the left two membranes. To corroborate this, a COMSOL model using the two left-most membranes as an areal concentration source of analyte was run. A horizontal slice of the computed concentration profile (S7f) as well as the linear and surface concentration gradients at the level of the sensor (S7h, i, respectively) both correlate very well with the experimental data shown in Figure S7e. A GIF file of the concentration profile over the first 500 seconds is shown in Figure S8.

### Sensing of Analytes through Disrupted Tissue Barrier

Finally, we tested sensor response to a large analyte (CRP) before and after selective HBE tissue barrier using TJDPs to demonstrate the utility of this platform for measuring barrier integrity and paracellular analyte transport in addition to cellular secretion. Development of a tissue barrier in vitro takes several days, so functionalized devices need to be able to maintain their functionality for days under flow in a 37 °C environment. To test this, single-channel devices were tested after 0, 4, and 6 days in the incubator. The results are shown in Figure S9, and confirm no loss of sensor functionality after 6 days. In previous work,^43^ we have demonstrated that TJDPs based on the amino acid sequence of Claudin 1 reversibly reduce barrier in HBEs, facilitating paracellular passage of large molecules in cell culture. Here, we first flowed CRP for 1 hour in the top channel of the device, and observed a modest 4 pm shift from the sensor in the bottom channel (Figure 8). This is consistent with the inability of the large CRP pentamer to permeate the barrier, which has formed tight junctions over the course of 4 days. Next, a TJDP was flowed at a concentration of 10 mM overnight, disrupting the barrier. After TJDP treatment, the sensors showed an 18 pm shift. As a positive control, CRP was briefly flowed in the bottom channel to directly address the sensors, yielding a 49 pm shift. Thus, a significant portion of the 1 μg/mL of CRP in the top channel was able to reach the sensors.

**Figure 8.**
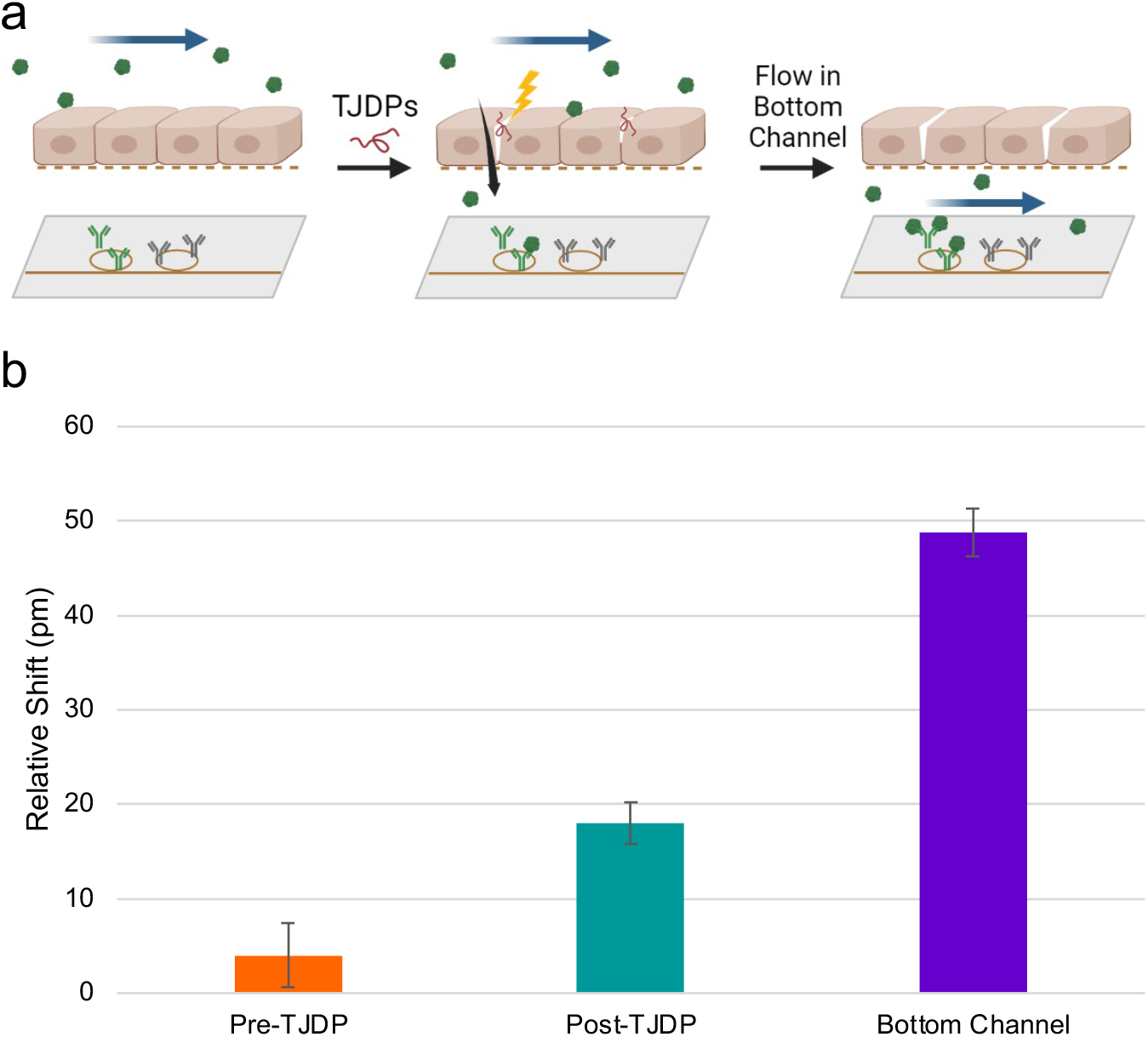
Barrier Disruption Sensing. a) Experimental diagram: CRP is flowed in the top channel before and after disruption with TJDPs, and sensed in the bottom channel. Finally, CRP is flowed directly into the bottom channel to confirm antibody reactivity. b) Quantification of shifts seen in each segment of the experiment. Error bars represent standard error of the mean, n = 7.

## Discussion

Existing in vivo and in vitro disease models have recognized shortcomings in their ability to facilitate understanding of human disease and predict therapeutic response^61^. The genetic heterogeneity of human clinical studies, the challenges of translating results of animal studies to humans, and the lack of temporally resolved information of appropriate biological complexity provided by in vitro cell culture systems all drive the need for new TC devices using human cells. Given the possibility of constructing such systems from iPSCs derived from an individual patient, TCs have immense promise in enabling patient-centric analysis of therapeutics, also known as the “n of 1 clinical trial”^62^. For TCs to reach their full potential, methods must be available to provide real-time data on the TC via incorporation of biosensors in close proximity to the cells under study. Photonic ring resonator biosensors are ideal for this application for several reasons, including their sensitivity, label-free nature, small footprint/large multiplexing capability, and scalable manufacture. Hence, we have integrated photonic ring resonators into a microfluidic system to sense the secretion of analytes from cells in real time.

The data described above confirms that our device design allows for culturing cells over the course of days, and performing experiments while attached to an optical setup used for sensing. Our photonic chips are able to sense physiologically relevant concentrations of analyte, reporting on the quantity of cytokines secreted by inflamed epithelial cells in real time following LPS stimulation. We also demonstrate detection of CRP, a large (120 kDa pentamer) protein passing through a disrupted cellular barrier. A particularly intriguing result is that these sensors may also be used to provide spatially-as well as temporally resolved information. These results provide proof of concept for incorporating photonic sensors into a broad range of more complex TC systems.

While these results represent a significant step forward in the development of tissue chip technology, there are a few limitations to our platform as currently constructed. In particular, the photonic sensors are able to quickly sense an increase in analyte concentrations, but not a decrease. This is due to the low off-rates of the antibodies with which the sensor is functionalized. One could envision using regeneration solutions to remove proteins from their corresponding antibodies, returning the sensor response to baseline. However, most common regeneration protocols involve extremes of pH or high salt concentrations^63,64^ that would lead to quick death of the TC under study. Thus, new regeneration protocols are needed that are safe for cells, or more sophisticated microfluidics that prevent regeneration solutions from coming into direct contact with the cells are required. Alternatively, modular systems in which new sensors are swapped into the microfluidic circuit when one set becomes saturated can also be envisioned.

The work presented here opens new opportunities for the study of physiological processes in TCs. While the inflammatory responses and conditional through-barrier transport measured here serve as useful proof-of-concept, the potential for our platform to learn new biology is great. The NPN membranes used as a culture substrate have excellent potential for various tissue barriers and the diseases that alter them, such as intestinal endothelium (Crohn’s disease^65^), skin (atopic dermatitis^66^), or the blood-brain barrier (Alzheimer’s^67^, traumatic brain injury^68,69^). All of these tissue barriers have multiple cell types that secrete paracrine markers that affect neighboring cell types. We anticipate that our new sensor-integrated platform can be used to determine the exact kinetics of protein release from varying cell types, leading to new understanding of disease pathophysiology. To further increase the kinds of data available from one of these experiments, sensors could also be situated in the top channel to enable assessment of the apical/basal secretion dynamics of tissue barriers.

Of course, this approach should also be useful in drug discovery and development. Currently, many drugs targeting the brain fail in clinical trials because they are unable to cross the blood-brain barrier (BBB). A platform such as the photonic sensor-enabled tissue chip, applied to BBB, could not only determine whether biological therapeutics can cross the BBB, but also measure cellular response in real time. Additionally, with the rapidly developing field of stem cell technology, the platform could easily be equipped with patient-specific induced pluripotent stem cells (iPSCs), to ensure patient response to a given therapeutic, making drugs both safer and more effective.

## Acknowledgements

The authors thank Michael Osadciw, for providing the illustration in Figure 1, Michael Pomerantz for producing custom aluminum heated stages, Ethan Luta for acquiring SEM images, and Dr. Dean Johnson for help with modeling nanoporous membranes in COMSOL.

## Author contributions

Conceptualization: JSC, BLM. Methodology: JSC, BLM, JLM. Investigation: JSC. COMSOL modeling: JSC, MTM. Cell culture protocols: MGB, JSC. Photonic chip design: MRB, JSC. Data analysis script development: JDT. Supervision: BLM. Writing-original draft: JSC, BLM. Writing—review & editing: JSC, BLM, JLM.

## Data and materials availability

All data as well as software used for the analysis of spectra are available from the corresponding author on request.

## Supplementary Figures

**Figure S1.**
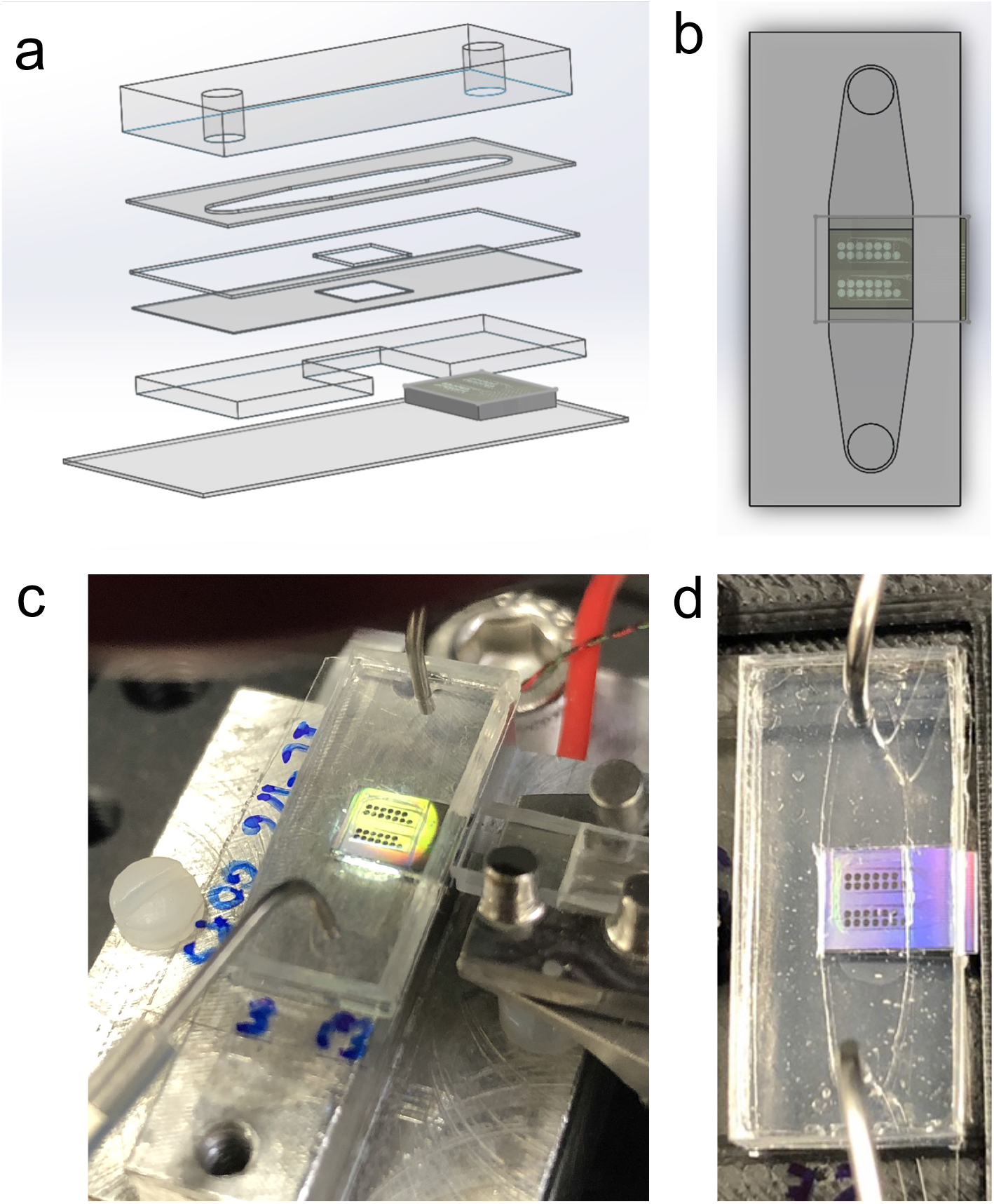
Single Channel Device. a) An exploded view of the device layers. A glass coverslip, silicone chip holder, two sealing layers, adhesive channel layer, and PDMS cap with inlet/outlet. b) Top view of the device with the photonic ring resonator sensors centered within the sealing layer and channel. c) Photo of the device on a temperature-controlled aluminum stage, addressed from the right by an 8-channel fiber array. d) Top view of the device under flow.

**Figure S2.**
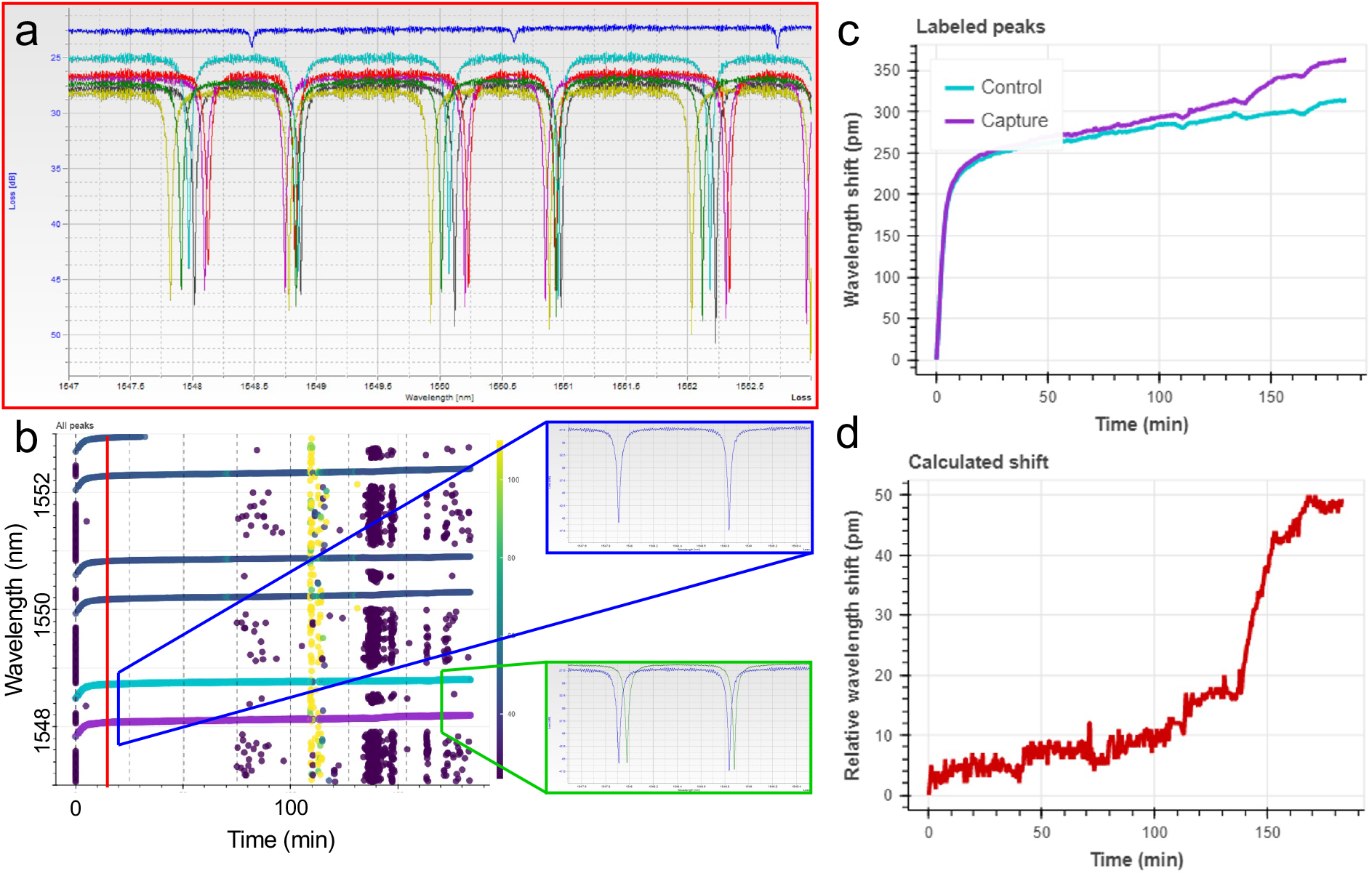
Data Analysis. a) Raw spectra for all 7 channels within Keysight’s viewer application. Each channel (distinguished by color) contains peaks for 2 rings, which are repeated approximately ever 2 nm, yielding 6 peaks in each 6-nm scan. b) Custom python analysis script user interface, which allows the user to select peaks at each timepoint and assign them to either the control or capture ring. Inset is an early timepoint (blue) and late timepoint (green), showing how the peaks move over time. Here the left/bottom peak is from the capture ring (IL-6 in the shown data) and in the spectrum from the later timepoint shows the left peak has shifted more than the right peak. c) A normalized plot of each peak’s location over time, showing a greater shift for the capture ring. d) Subtracted shift, showing the relative wavelength shift of the capture ring vs. the control ring. This data shows a calibration of IL-6. The x-axis represents time but roughly corresponds to concentration, with the final shift of about 48 pm occurring at the highest concentration of IL-6 (1 μg/mL).

**Figure S3.**
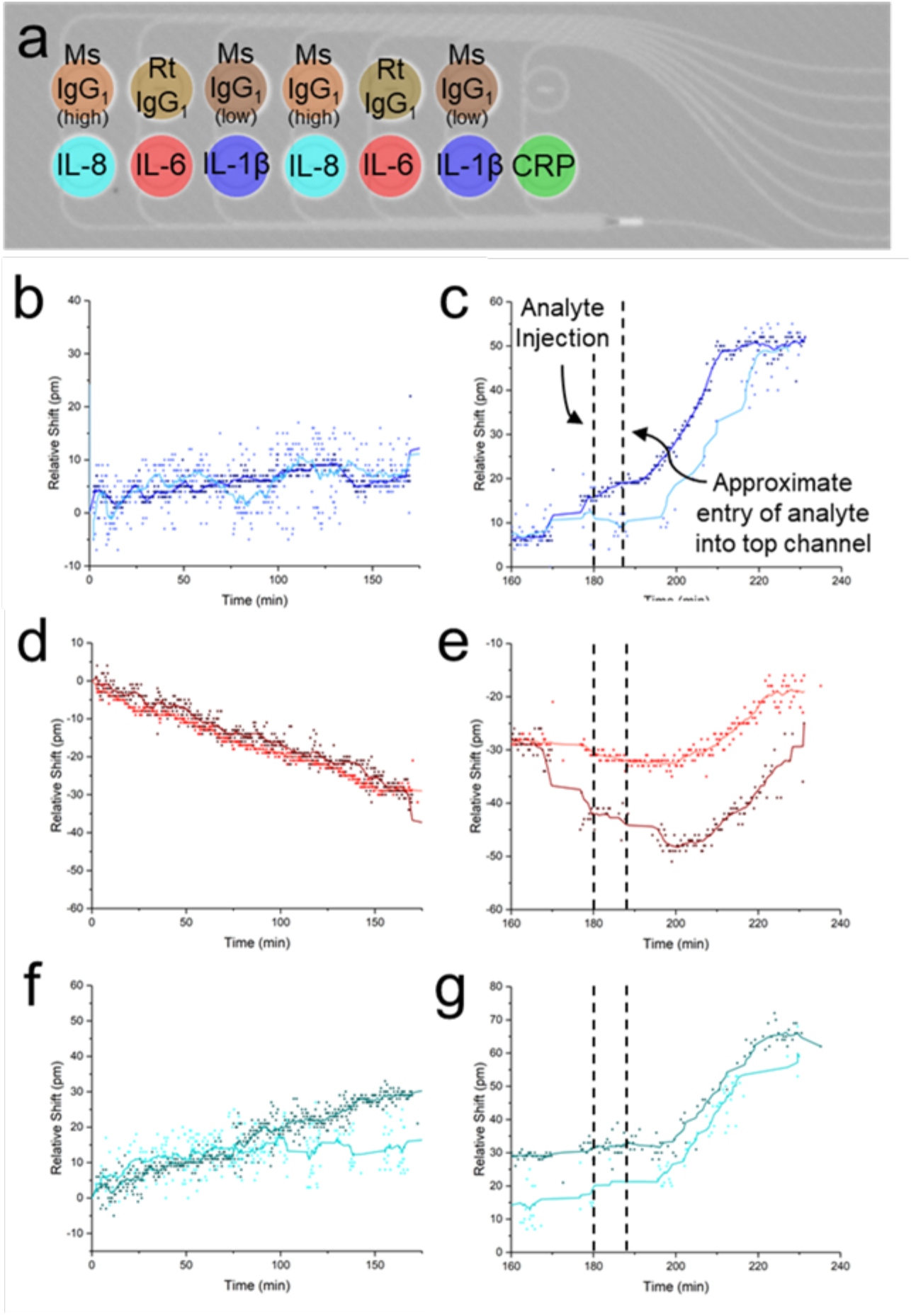
Nonspecific Binding and Diffusion Through Bottom Channel. a) Ring functionalization scheme. Each control antibody is matched to the antibody isotype of the corresponding capture antibody. “High” and “Low” for the mouse antibodies refers to the concentration used to match that of IL-1β vs IL-8, since their stock concentrations were different. b) Nonspecific binding for two IL-1β channels (both represent control-subtracted relative shifts) over about 3 hours, and c) relative shifts after adding analyte. d) a significant relative blueshift is seen for IL-6, meaning the isotype control antibody used is not an ideal match. e) Significant redshifts seen for IL-6 once analyte is added. f) Nonspecific and g) analyte-specific shifts for IL-8.

**Figure S4.**
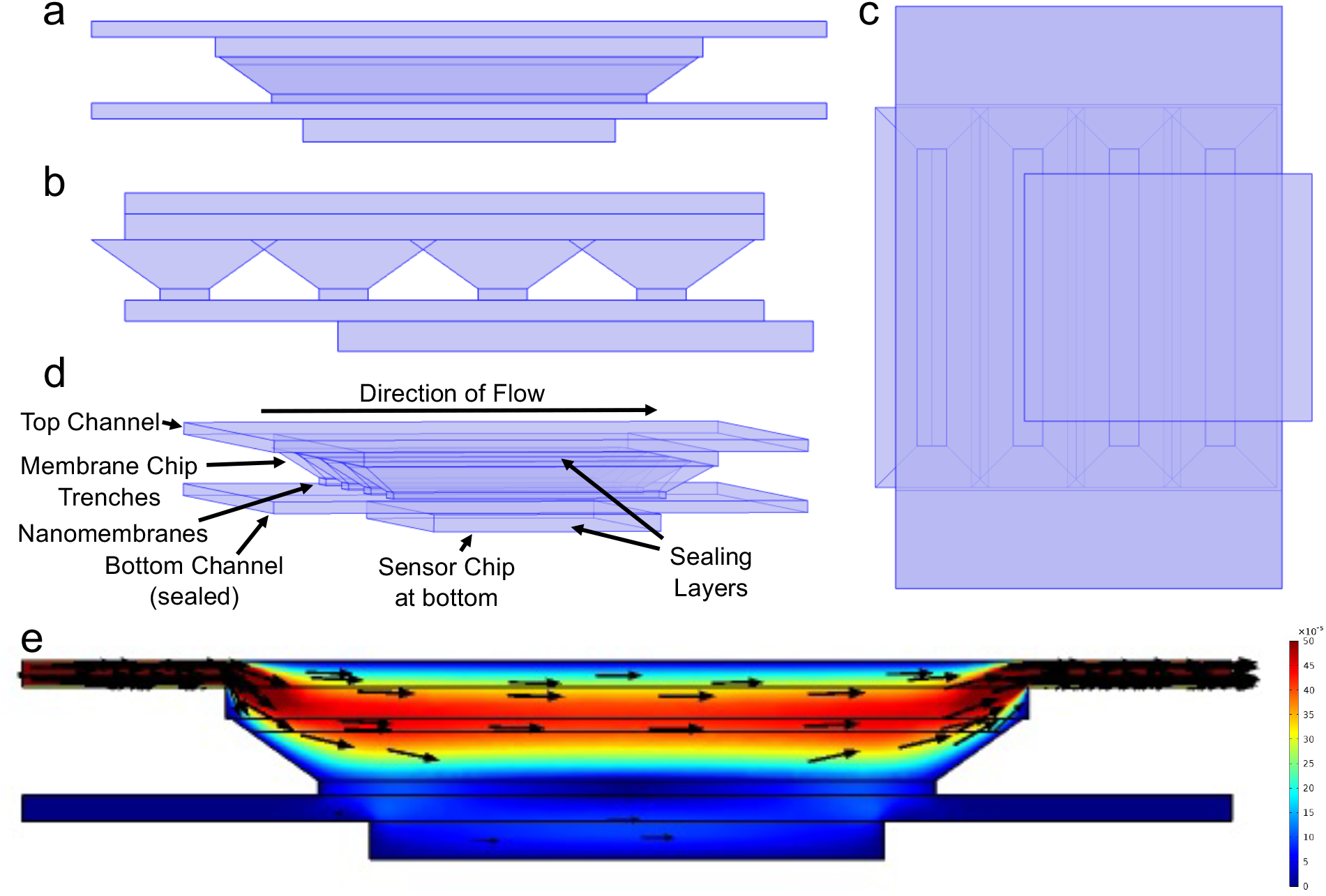
COMSOL Fluid Flow Model. a) Side view, b) end view, c) top view and d) oblique view of the device geometry used for COMSOL simulations. Top and bottom channels sandwich the four membranes, which have a trapezoidal trench above them, sealed with silicone and adhesive on top. The photonic chip is below the bottom channel and a sealing layer. e) Fluid flow simulation with a constant volumetric flow rate of 30 μL/min in the top channel, with the bottom channel sealed (streamlines are proportional in length to velocity). The membranes effectively isolate the bottom channel from flow, resulting in largely diffusion-mediated transport.

**Figure S5.**
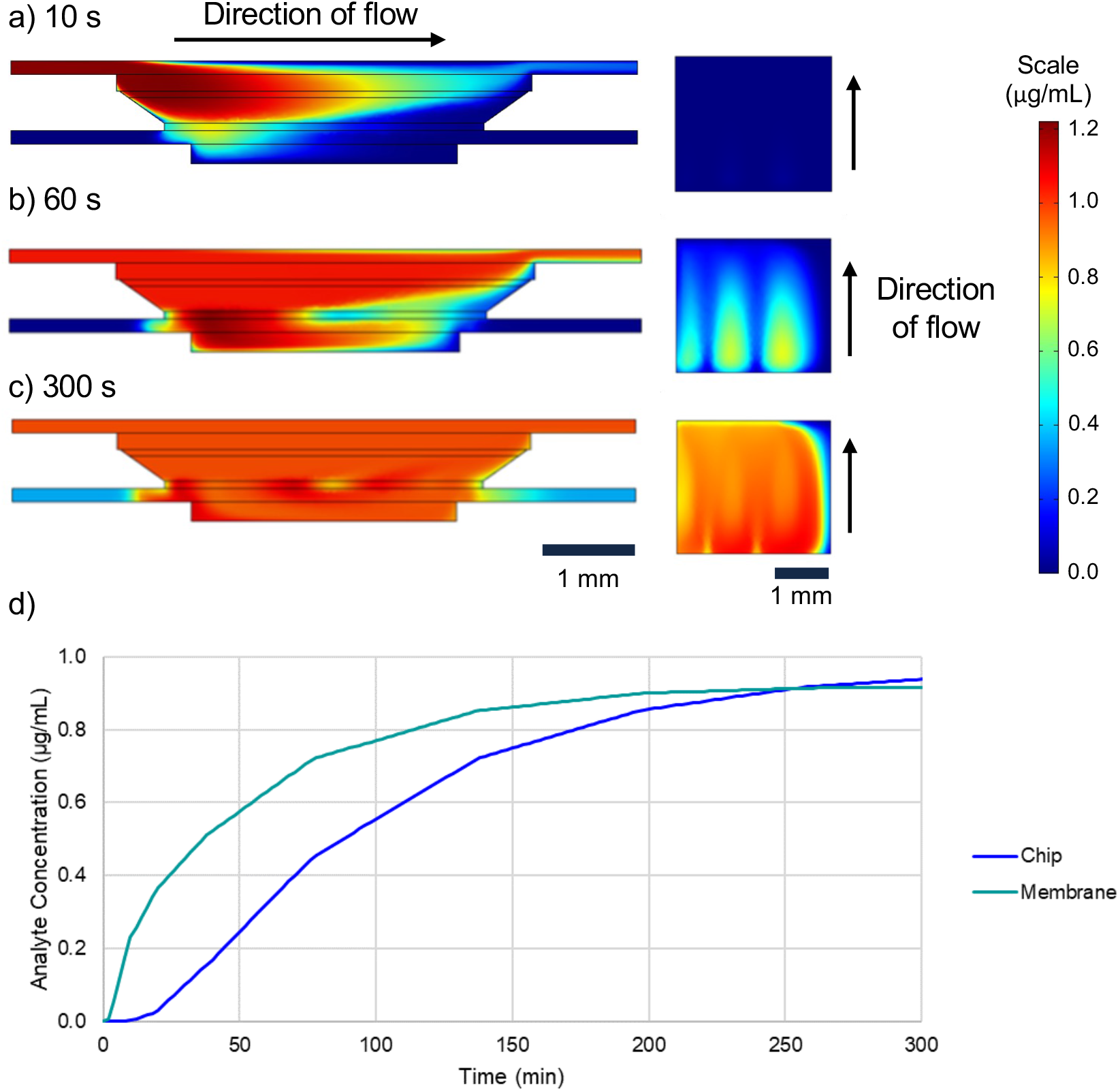
COMSOL Concentration Profile. Side view concentration profile at a) 10 s, b) 60 s, and c) 300 s. The left image is a side view of the device with flow in the top channel going left to right. The square on the right is a top view of the sensor surface, with rectangles corresponding with the three membranes that are situated over the chip. d) Surface average of the concentration at the level of the membrane (teal line) and sensor chip (blue) over time.

**Figure S6.**
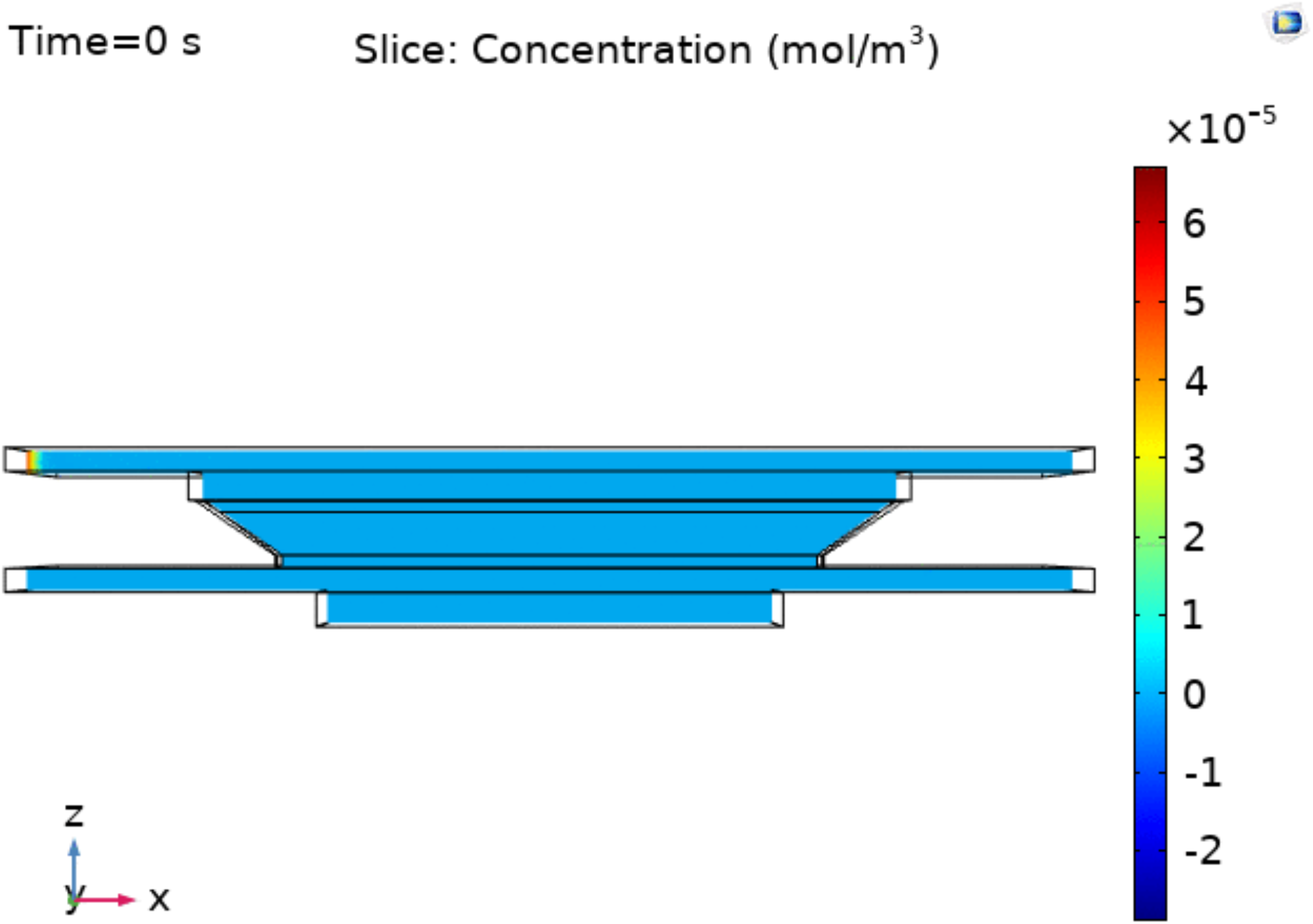
Concentration Profile Time Course. A gif file showing the concentration of IL-6 in the device as a function of time, up to 500 s. Here a constant source of analyte (1 μg/mL) is flowed in the top channel at 30 μL/min and the bottom channel is static to allow for diffusion to the level of the sensor surface. The scale bar is in COMSOL default units, with 5E-5 corresponding to 1 μg/mL.

**Figure S7.**
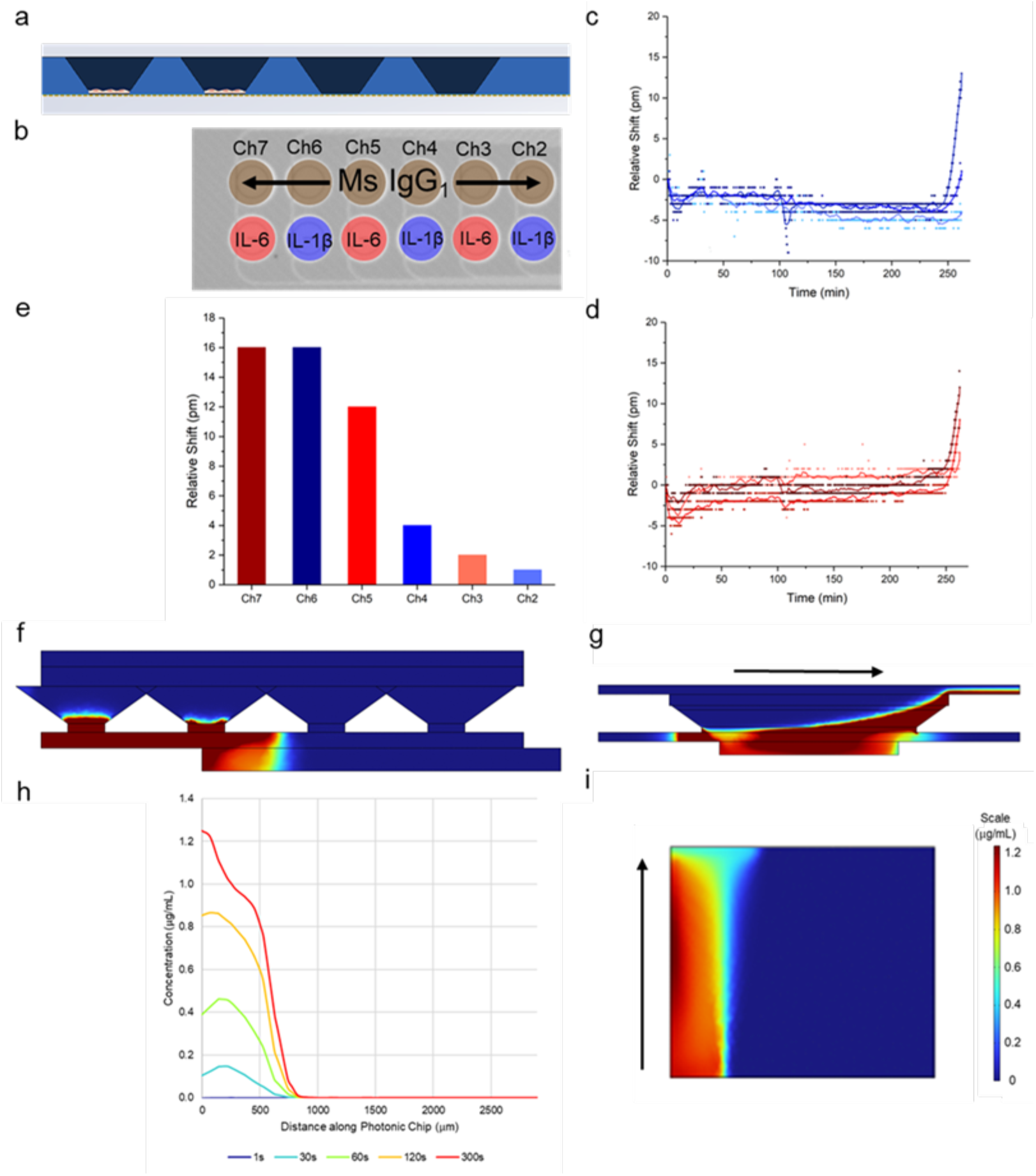
Partial Cell Layer LPS Stimulation Experiment. a) Diagram of the nanomembrane chip with cells removed from the two right-most membranes. b) Photonic chip functionalization schematic with alternating IL-6- and IL-1β-functionalized channels. The furthest-right edge of the source of cellular secretion approximately lines up with Channel 7. c) Subtracted traces for channels 2, 4, and 6 (IL-1B) and d) 3, 5, and 7 (IL-6) over time. e) Maximum relative shift for each channel moving left to right across the photonic chip. f) A COMSOL simulation showing the cross section of the channel showing the diffusion of IL-6 in the bottom channel after 5 minutes. g) A side view of the same simulation, showing that while analyte secreted into the top channel, which has a flow rate of 30 uL/min, is immediately swept out via the top outlet, the sealed bottom channel allows the analyte to diffuse to the level of the sensor. h) Quantification of IL-6 concentration across the width of the photonic chip at different time points. i) Concentration of IL-6 on the surface of the photonic chip, with the analyte flowing bottom-to-top in the top channel (though there is no flow directly above the chip). Panels a, b, e, f, and h are aligned vertically to show the relation between the membranes, sensors, and cytokine concentrations across the device cross-section.

**Figure S8.**
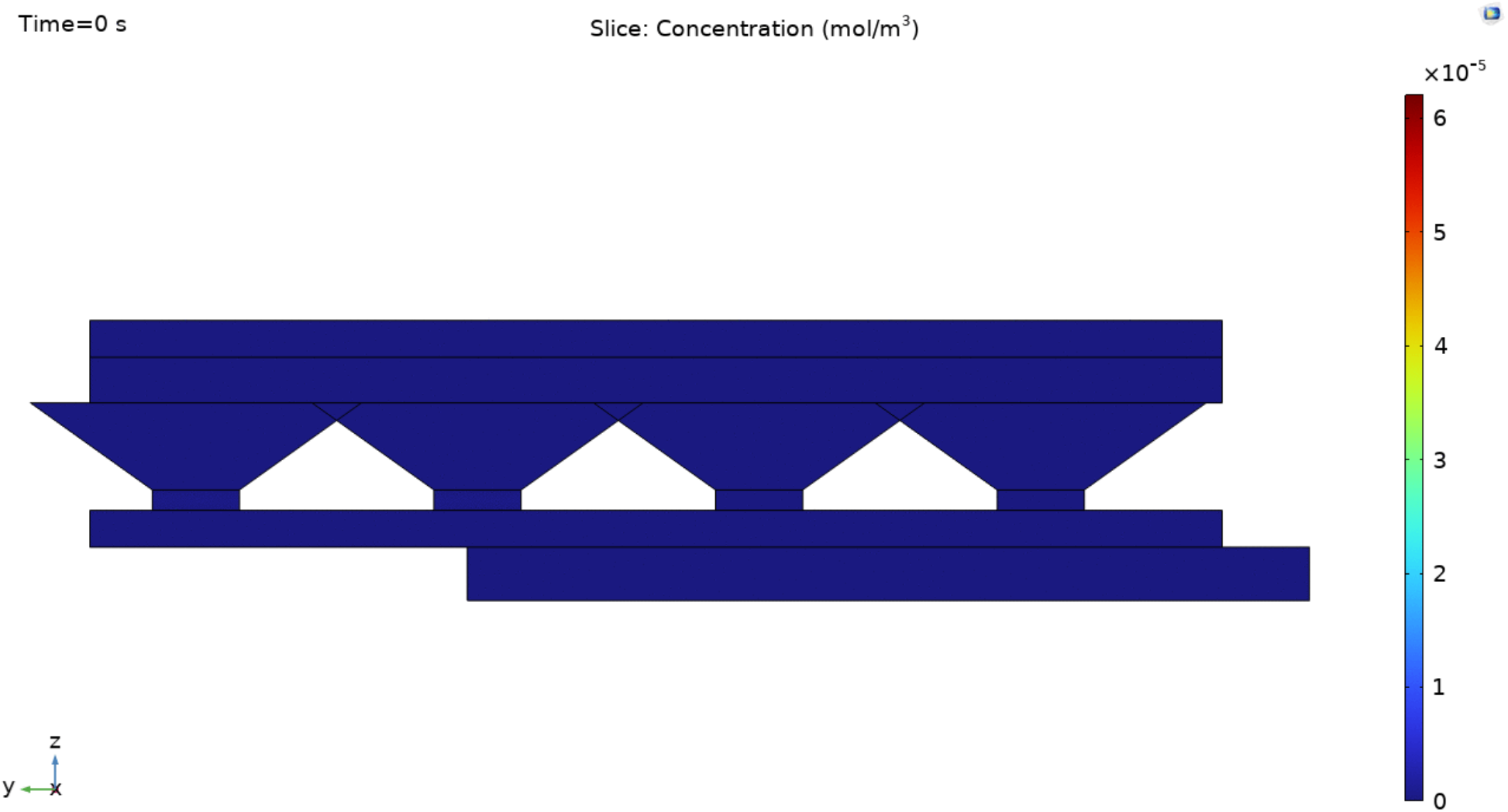
Partial Cell Layer Concentration Profile Time Course. A gif file showing the diffusion of IL-6 in a 2-channel microfluidic device with sources of cellular secretion (1 μg/mL) at the two left-most membranes.

**Figure S9.**
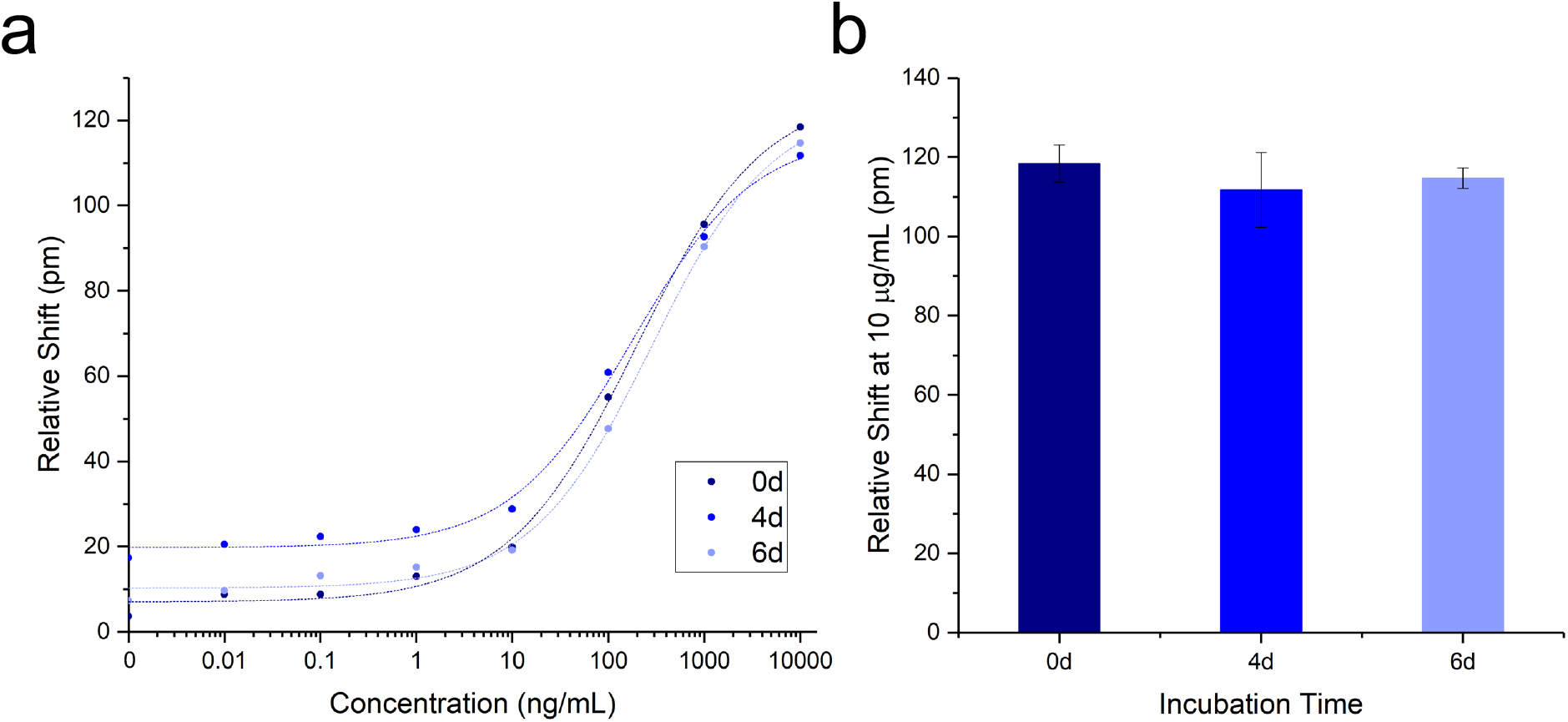
Antibody Stability in Incubator. a) Concentration curves for CRP in single-channel microfluidic devices, incubated at 37°C for 0, 4, and 6 days. b) Maximum shift (at 10 μg/mL) for each timepoint shows negligible effect of incubation on antibody affinity.

