## Supplementary figures and images for "A Photonic Biosensor-Integrated Tissue Chip Platform for Real-Time Sensing of Lung Epithelial Inflammatory Markers"

### Supplementary Figure S6

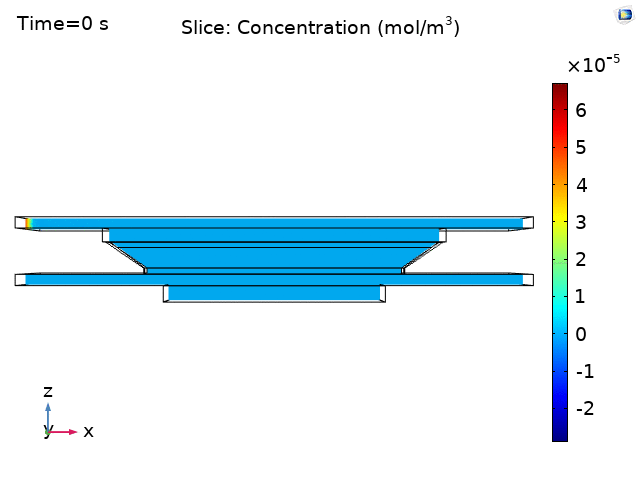

### Supplementary Figure S8

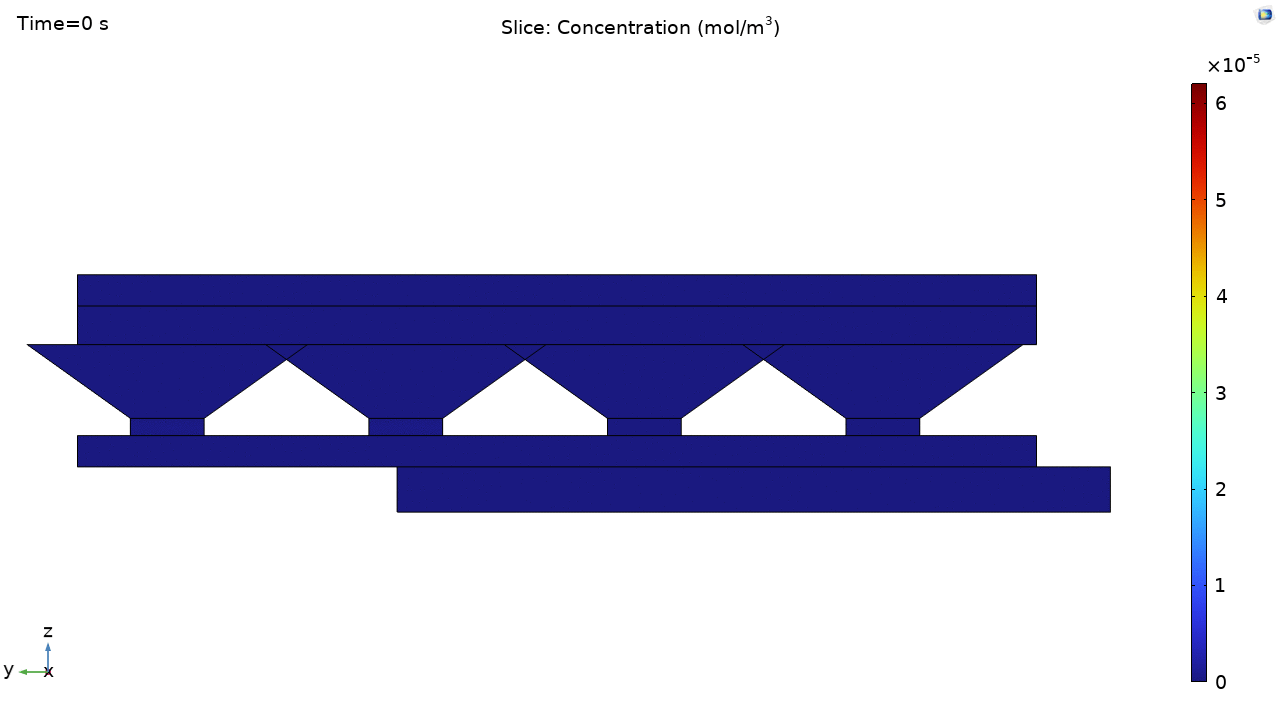
